# Brain white matter pathways of resilience to chronic back pain: a multisite validation

**DOI:** 10.1101/2024.01.30.578024

**Authors:** Mina Mišić, Noah Lee, Francesca Zidda, Kyungjin Sohn, Katrin Usai, Martin Löffler, Md Nasir Uddin, Arsalan Farooqi, Giovanni Schifitto, Zhengwu Zhang, Frauke Nees, Paul Geha, Herta Flor

## Abstract

Chronic back pain (CBP) is a global health concern with significant societal and economic burden. While various predictors of back pain chronicity have been proposed, including demographic and psychosocial factors, neuroimaging studies have pointed to brain characteristics as predictors of CBP. However, large-scale, multisite validation of these predictors is currently lacking. In two independent longitudinal studies, we examined white matter diffusion imaging data and pain characteristics in patients with subacute back pain (SBP) over six- and 12-month periods. Diffusion data from individuals with CBP and healthy controls (HC) were analyzed for comparison. Whole-brain tract-based spatial statistics analyses revealed that a cluster in the right superior longitudinal fasciculus (SLF) tract had larger fractional anisotropy (FA) values in patients who recovered (SBPr) compared to those with persistent pain (SBPp), and predicted changes in pain severity. The SLF FA values accurately classified patients at baseline and follow-up in a third publicly available dataset (Area under the Receiver Operating Curve ∼ 0.70). Notably, patients who recovered had FA values larger than those of HC suggesting a potential role of SLF integrity in resilience to CBP. Structural connectivity-based models also classified SBPp and SBPr patients from the three data sets (validation accuracy 67%). Our results validate the right SLF as a robust predictor of CBP development, with potential for clinical translation. Cognitive and behavioral processes dependent on the right SLF, such as proprioception and visuospatial attention, should be analyzed in subacute stages as they could prove important for back pain chronicity.

## Introduction

Chronic back pain (CBP) is the leading cause of disability worldwide(1); the annual contribution of patients with back pain to all-cause medical costs is $365 billion in the United States alone.(2) In 2020, there were over 500 million reported cases of prevalent low back pain globally and central Europe had the highest age-standardized rate of prevalence with 12,800 cases per 100,000 individuals.(3) In addition, CBP is associated with significant co-morbidities such as substance misuse,(4) depression,(5–7) anxiety,(5, 7) and obesity.(8, 9) The large majority of CBP cases suffer from primary pain.(10) It is estimated that 25-50 % of patients with subacute back pain (SBP, duration 6-12 weeks) go on to develop CBP.(11–15) Once back pain is chronic, it is very difficult to treat.(16–20) Therefore, prevention of the transition to the chronic phase(21) is key to reducing the prevalence, and as a result, the burden of CBP.

Early identification of the likelihood of pain chronicity is a pre-requisite step for allowing the efficient implementation of preventive treatment (22) because diagnostic and treatment procedures offered to all (sub)acute back pain patients expose resilient patients to unnecessary and risky procedures. In addition, treatment offered to all patients is burdensome to the healthcare system.(23, 24) However, validated quantitative prognostic biomarkers that can predict disease trajectory and guide treatment in back pain patients have yet to be identified.

Some studies have addressed this question with prognostic models incorporating demographic, pain-related, and psychosocial predictors.(21, 25–27) While these models are of great value showing that few of these variables (e.g. work factors) might have significant prognostic power on the long-term outcome of back pain, their prognostic accuracy is limited,(28) with parameters often explaining no more than 30% of the variance.(29–31) A recent notable study in this regard developed a model based on easy-to-use brief questionnaires to predict the development and spread of chronic pain in a variety of pain conditions capitalizing on a large dataset obtained from the UK-BioBank. (32) This work demonstrated that only a few features related to the assessment of sleep, neuroticism, mood, stress, and body mass index were enough to predict persistence and spread of pain with an area under the curve of 0.53-0.73. Yet, this study is unique in showing such a predictive value of questionnaire-based tools. Neurobiological measures could therefore complement existing prognostic models based on psychosocial variables to improve overall accuracy and discriminative power. More importantly, neurobiological factors such as brain parameters can provide a mechanistic understanding of chronicity and its central processing.

Neuroimaging research on chronic pain has uncovered a shift in brain responses to pain when acute and chronic pain are compared. The thalamus, primary somatosensory, motor areas, insula, and mid-cingulate cortex most often respond to acute pain and can predict the perception of acute pain.(33–36) Conversely, limbic brain areas are more frequently engaged when patients report the intensity of their clinical pain.(37, 38) Consistent findings have demonstrated that increased prefrontal-limbic functional connectivity during episodes of heightened subacute ongoing back pain or during a reward learning task is a significant predictor of CBP.(39, 40) Furthermore, low somatosensory cortex excitability in the acute stage of low back pain was identified as a predictor of CBP chronicity.(41) One study so far has investigated structural changes in back pain using a longitudinal design and found that changes across several white matter tracts including the temporal part of the left superior longitudinal fasciculus, external capsule, parts of the corpus callosum, and parts of the internal capsule were predictive of back pain development at 1-year follow-up.(42) Despite the identification of these promising biomarkers of pain chronicity, their validation across multiple independent cohorts remains rare.(43) Here we investigated brain white matter predictors of back pain chronicity across three independent samples originating from 3 different sites and expected white matter tracts previously found predictive of CBP(42) to show higher fractional anisotropy values (FA; greater structural integrity) in patients who recovered compared to those whose pain persisted. We additionally hypothesized that patients who recovered would also exhibit greater structural integrity within the prefrontal-limbic tract related to learning and memory, uncinate fasciculus,(44) in line with findings that learning-related processes could be predictive of CBP.(40) We expected that lower FA at baseline would predict persistent state and greater pain severity at follow-up.

## Results

### Demographic and clinical characteristics

At baseline, the data collected in New Haven included 16 SBP patients who recovered (SBPr) at approximately one-year follow-up as their low-back pain intensity dropped by more than 30% relative to baseline and 12 SBP patients whose pain persisted at follow-up (**Table S1**). The SBPp patients were older (38.0 ± 3.6 years, average ± SEM) and had pain for a slightly longer duration (10.8 ± 0.9 weeks) than the SBPr patients (age = 30.8 ± 2.2 years; pain duration = 8.6 ± 0.9 weeks) but these differences did not reach statistical significance (t-score (degrees of freedom) (*t*(df)) = 1.8(26), *p* = 0.08 for age and *t*(df) =1.8(26), *p* = 0.08 for duration comparisons, respectively, unpaired T-test). The groups did not significantly differ in the distribution of males and females (𝜒2= 0.32, df = 1, *p* = 0.57) or body mass index (BMI) (*t*(df) = 0.35(26), *p* = 0.73). They did not significantly differ in average reported pain intensity (*t*(df) = - 0.45(26), *p* = 0.66) (**Table S1**). The SBPr patients reported significantly (*p* = 0.02) larger depression scores on Beck’s Depression Inventory (BDI) (7.3 ± 1.2) than the SBPp patients (3.1 ± 1.1, *t*(df) = - 2.65(26), *p* = 0.02) but the average score indicated that the SBPr patients did not have any clinically significant symptoms (i.e., average BDI < 10).

### Whole-brain Tract-Based Spatial Statistics

We calculated voxel-wise FA of each participant and performed a whole brain comparison over the white matter skeleton using permutation testing between SBPr (*n* = 16) and SBPp (*n* = 12) patients at baseline (unpaired t-test, *p* < 0.05, cluster-based thresholding) corrected for age, gender, and head displacement estimated by eddy current correction (**Fig. S2**). SBPr showed larger FA within the cluster of fibers part of the right superior longitudinal fasciculus (SLF) (MNI-coordinates of peak voxel: x = 35; y = - 13; z = 26 mm; t(max) = 4.61) (**Fig. 1**). The possibility that the observed difference between SBPr and SBPp patients was due to a difference in the amount of displacement applied during registration was ruled out (**Fig. S2**). We plotted the FA values of the same SLF region from healthy controls (HC) and CBP patients; **Fig. 1** shows FA values and their distributions for each group within the SLF cluster. SBPr patients had the largest FA values in the right SLF cluster, even larger than in HC, although this difference did not reach statistical significance (*p* = 0.11). We also examined mean diffusivity (MD), axial diffusivity (AD), and radial diffusivity (RD) extracted from the right SLF shown in **Fig.1** to further understand which diffusion component is different between the groups. The right SLF MD is significantly increased (p < 0.05) in the SBPr compared to SBPp patients (**Fig. S3**), while the right SLF RD is significantly decreased (*p* < 0.05) in the SBPr compared to SBPp patients in the New Haven data (**Fig. S4**). Axial diffusivity extracted from the RSLF mask did not show significant difference between SBPr and SBPp (*p* = 0.28) (**Fig. S5**).

**Figure 1.**
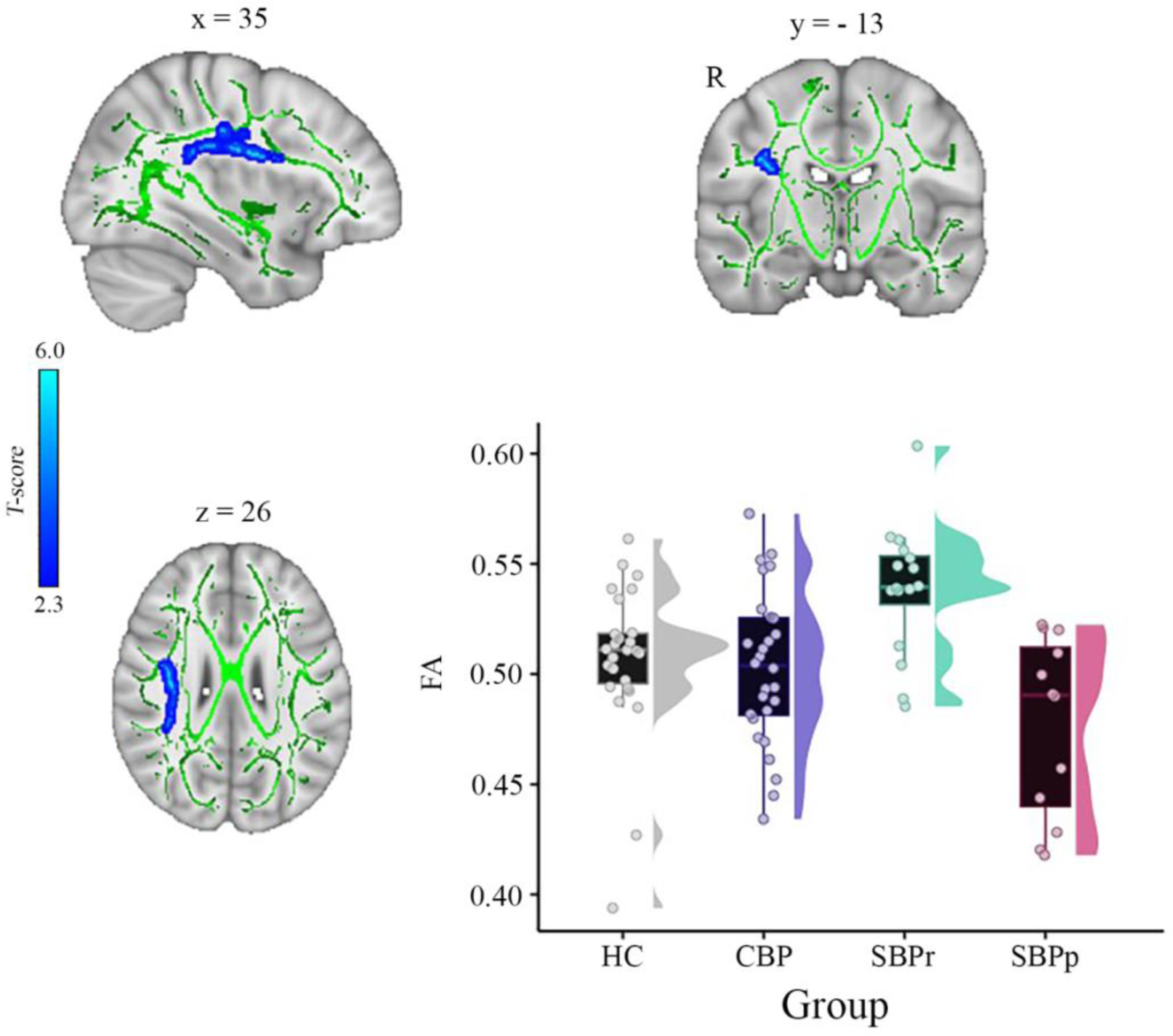
A whole brain comparison over the white matter skeleton between SBPr and SBPp patients at baseline and distribution of FA values for each group (New Haven data set). Results of unpaired t-test (p < 0.05, 10,000 permutations) showing significantly increased FA (fractional anisotropy) in SBPr (recovered) compared to SBPp (persistent) patients within the right superior longitudinal fasciculus (SLF) in the New Haven data set. Rain clouds include boxplots and the FA data distribution for each group depicted on the right side of each boxplot. Jittered circles represent single data points, the middle line represents the median, the hinges of the boxplot the first and third quartiles, and the upper and lower whiskers 1.5*IQR (the interquartile range).

To test whether baseline FA values predict the change in pain severity from baseline to follow-up in the New Haven data set, we did multiple regression analysis with FA values as predictor and the percentage change in pain severity as outcome. In this model, FA values were predictive of pain severity at the one-year follow-up (adjusted 𝑅^2^ = 0.202, *p* = 0.009). To confirm that this result was not driven by age, gender, or head motion, we entered these parameters in a new model adjusting the prediction for these covariates. FA values were still predictive of the change in pain intensity, with added variables improving the model fit (new model: adjusted 𝑅^2^ = 0.259, *p* = 0.037; difference between models: *F*(2,67) = 1.50, *p* = 0.237). **Fig. 2** depicts the correlation between FA values in the right SLF and pain severity with higher FA values (greater structural integrity) associated with greater reduction in pain (percentage change).

**Figure 2.**
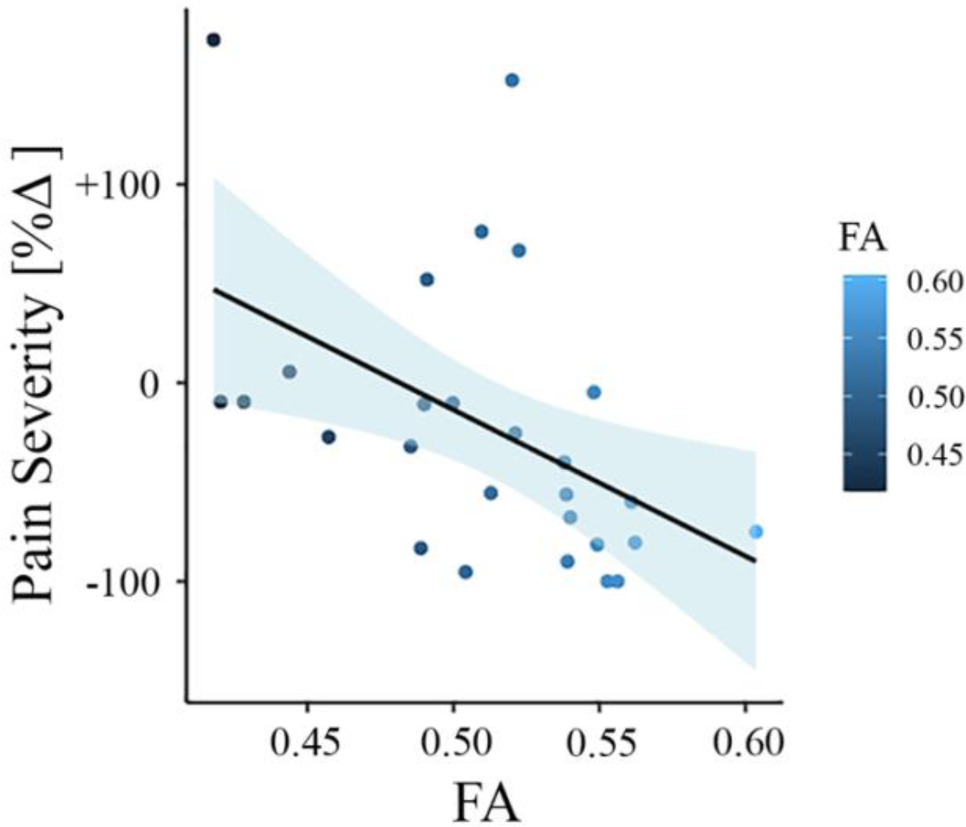
Association between white matter FA values and pain severity (New Haven data set). Higher fractional anisotropy (FA) values in the right superior longitudinal fasciculus (SLF) are associated with greater pain reduction (from baseline to follow-up) in the New Haven data set.

### Validation of the results obtained from the New Haven data

#### Mannheim data

In another independent study, a whole-brain comparison of FA over the white matter skeleton using permutation testing (unpaired t-test, *p* < 0.05, threshold-free cluster enhancement (TFCE) corrected for age, gender, and two motion parameters (translation and rotation) revealed two clusters, one in the right superior longitudinal fasciculus (SLF) tract (cluster size = 409 voxels, MNI-coordinates of peak voxel: x = 26, y = −33, z = 45, *p*(TFCE) = 0.041, t(max) = 3.57) and one in the right corticospinal tract/superior corona radiata (cluster size = 381 voxels, MNI-coordinates of peak voxel: x = 29, y = −16, z = 21, *p*(TFCE) = 0.041, t(max) = 3.63) that were significantly greater in SBPr (*N* = 28) compared to SBPp (*N* = 18) (**Fig. 3**). In addition, two smaller clusters were also identified in the same tract of the right SLF (cluster size = 39 voxels, MNI-coordinates of peak voxel: x = 36, y = −13, z = 34, *p*(TFCE) = 0.048, t(max) = 3.19) and right corticospinal tract (cluster size = 13 voxels, MNI-coordinates of peak voxel: x = 21, y = −27, z = 42, *p*(TFCE) = 0.049, t(max) = 2.41) as significantly different between SBPr and SBPp patients. In the next step, we extracted FA values from the significant clusters of the Mannheim data and compared them across all groups at that site. As in the New Haven set, recovered patients had the largest FA values across groups, even greater than HC although this difference did not reach significance level (*p* = 0.12) (**Fig. 3**).

**Figure 3.**
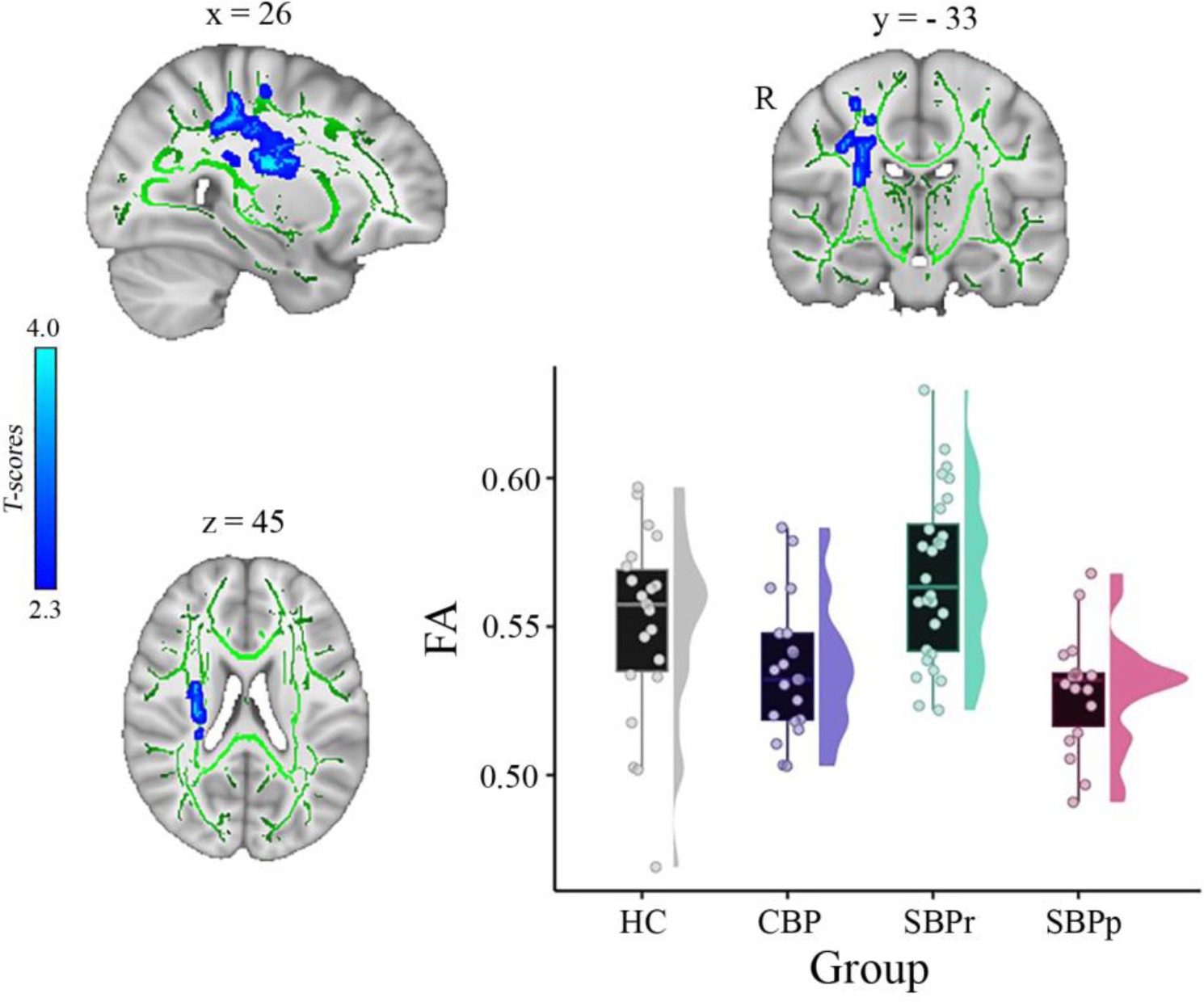
A whole brain comparison over the white matter skeleton between SBPr and SBPp patients at baseline and distribution of FA values for each group (*Mannheim data set*). Results of unpaired t-test (p < 0.05, 10,000 permutations) showing significantly increased FA (fractional anisotropy) in SBPr (recovered) compared to SBPp (persistent) patients at six-months follow-up within the right superior longitudinal fasciculus (SLF) in the Mannheim data set. Rain clouds include boxplots and the FA data distribution for each group depicted on the right side of each boxplot. Jittered circles represent single data points, the middle line represents the median, the hinges of the boxplot the first and third quartiles, and the upper and lower whiskers 1.5*IQR (the interquartile range).

To test whether FA baseline values from the significantly different clusters could predict the change in pain severity from baseline to follow-up, we did multiple regression analysis with FA values as predictor and the change in pain severity percentage as outcome. In this model, FA values were predictive of pain severity at the 6-month follow-up (adjusted 𝑅^2^ = 0.120, *p* = 0.011). To confirm that this result was not driven by age, gender, or head motion, we entered these parameters in a new model adjusting the prediction for covariates. FA values were still predictive of chronicity, with added variables improving the model fit (new model: adjusted 𝑅^2^ = 0.236, *p* = 0.007; difference between models: *F*(2,67) = 3.67, *p* = 0.046). **Fig. 4** depicts the correlation between FA values and pain severity with higher FA values (greater structural integrity) associated with greater reduction in pain (percentage change).

**Figure 4.**
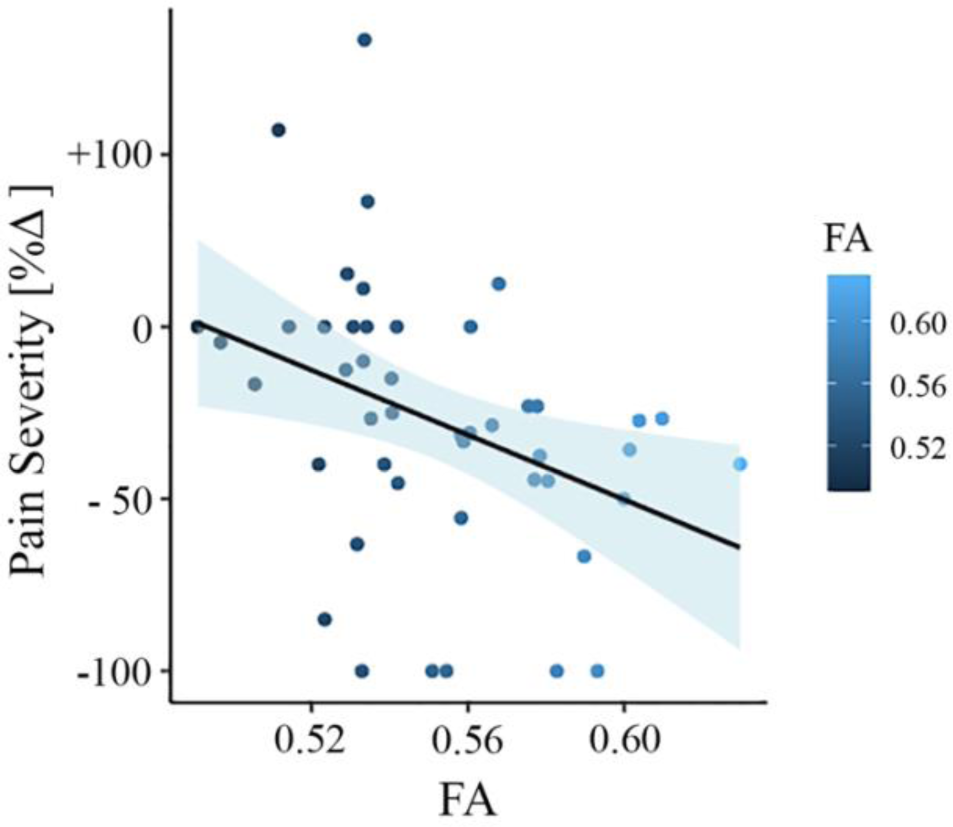
Association between white matter FA values and pain severity (Mannheim data set). Higher fractional anisotropy (FA) values in the right superior longitudinal fasciculus (SLF) are associated with greater pain reduction (from baseline to follow-up) in the Mannheim data set.

We tested the accuracy of local diffusion properties of the right SLF extracted from the mask of voxels passing threshold in the New Haven data (**Fig. 1**) in classifying the Mannheim patients into persistent and recovered. We used a simple cut-off(45) for the evaluation of the area under the receiver operating characteristic (ROC) curve (AUC). FA values corrected for age, gender, and head displacement, accurately classified SBPr (*N* = 28) and SBPp (*N* = 18) patients from the Mannheim data set with an AUC = 0.66 (*p* = 0.031, tested against 10,000 random permutations, see **Fig. S7A**), validating the predictive value of the right SLF cluster (**Fig. 5**).

**Figure 5.**
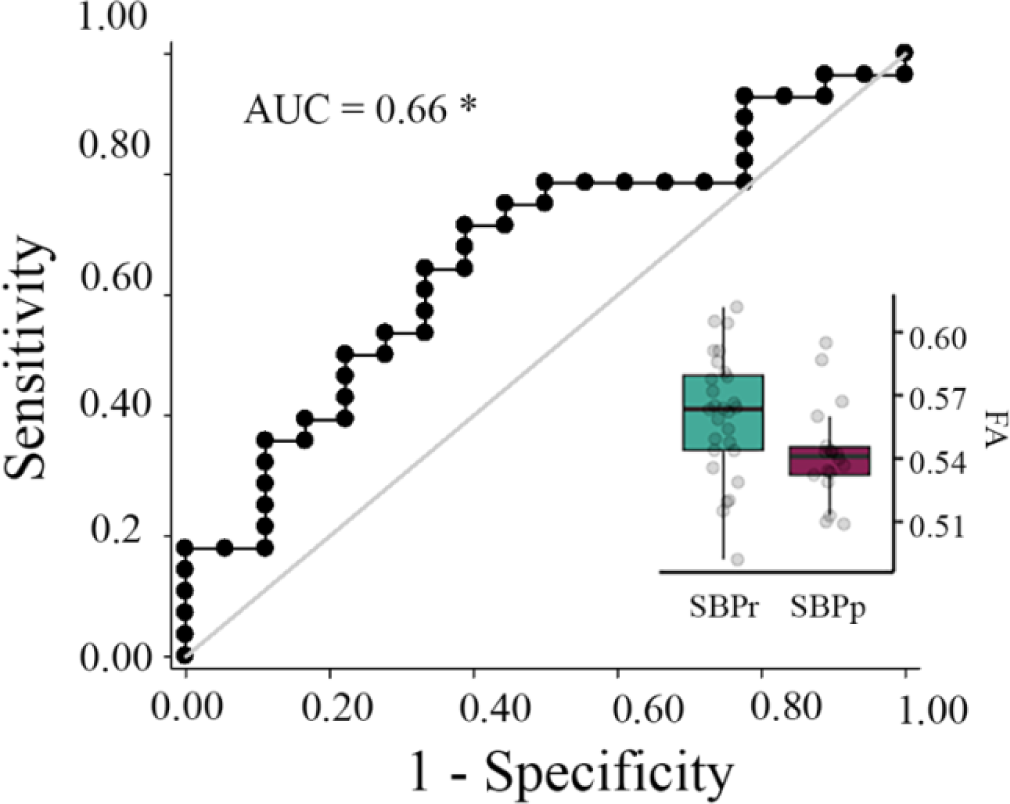
Validation of the accuracy of FA in the right SLF in classifying Mannheim patients. The right superior longitudinal fasciculus (SLF) cluster from the discovery set accurately classifies patients who recovered (SBPr) and those whose pain persisted (SBPp) in the Mannheim data set at a six-month follow-up. Classification accuracy is based on the ROC curve. Circles on the boxplots represent single data points, the middle line represents the median, the hinges of the boxplot the first and third quartiles, and the upper and lower whiskers 1.5*IQR (the interquartile range). AUC: area under the curve; * *p* < 0.05.

Supplementary Figure S6 shows the results in the Mannheim data set if a 30% reduction is used as a recovery criterion in this dataset (AUC= 0.53).

#### Chicago data

To further validate the right SLF predictive power, we used the mask shown in **Fig. 1** to extract FA values from SBPr and SBPp patients available in the Chicago data set. FA values in the validation data sets were corrected for age, gender, and head displacement. The criterion for recovery was set identically to the New Haven study, requiring ≥ 30% reduction of reported low-back pain intensity at one-year follow-up. FA values were calculated for two time points; one obtained at baseline when pain was still subacute (6-12 weeks), and one obtained at a one-year follow-up when pain either remitted or persisted.(46) FA values of the right SLF (**Fig. 1**) accurately classified SBPr (*N* = 23) and SBPp (*N* = 35) patients from Chicago with an AUC = 0.70 (*p* = 0.0043, **Fig. S7B**) at baseline (**Fig. 6A**), and SBPr (*N* = 28) and SBPp (*N* = 34) patients with an AUC = 0.66 (*p* = 0.014, see **Fig. S7C**) at follow-up (**Fig. 6B**), validating the predictive cluster from the right SLF at yet another site. The correlation between FA values in the right SLF and pain severity in the Chicago data set showed marginal significance (*p* = 0.055) at visit 1 (**Fig. S8A)** and higher FA values were significantly associated with a greater reduction in pain at visit 2 (*p* = 0.035) (**Fig. S8B**).

**Figure 6.**
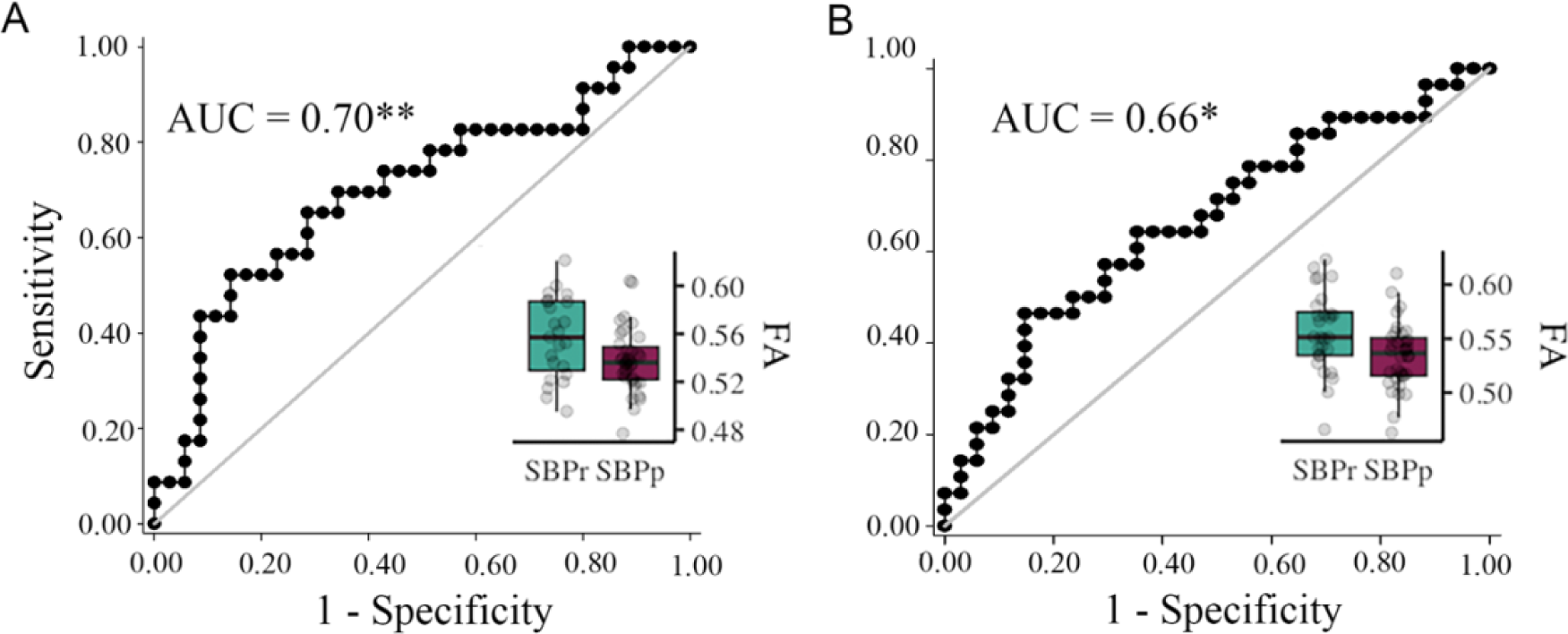
Validation of the accuracy of FA in the right SLF in classifying Chicago patients. The right SLF (superior longitudinal fasciculus) cluster from the discovery set accurately classifies patients who recovered (SBPr) and those whose pain persisted (SBPp) in the Chicago (OpenPain) data set at visit 1 (baseline) **(A)** and visit 2 (one-year follow-up) **(B)**. Classification accuracy is based on the ROC curve. Circles on the boxplots represent single data points, the middle line represents the median, the hinges of the boxplot the first and third quartiles, and the upper and lower whiskers 1.5*IQR (the interquartile range). AUC: area under the curve; * *p* < 0.05.

### Validation after harmonization

Because the DTI data sets originated from 3 sites with different MR acquisition parameters, we repeated our TBSS and validation analyses after correcting for variability arising from site differences using DTI data harmonization as implemented in neuroCombat. (47) The method of harmonization is described in detail in the Supplementary Methods. The whole brain unpaired t-test depicted in **Figure 1** was repeated after neuroCombat and yielded very similar results (**Fig. S9A**) showing significantly increased FA in the SBPr compared to SBPp patients in the right superior longitudinal fasciculus (MNI-coordinates of peak voxel: x = 40; y = - 42; z = 18 mm; t(max) = 2.52; p < 0.05, corrected against 10,000 permutations). We again tested the accuracy of local diffusion properties (FA) of the right SLF extracted from the mask of voxels passing threshold in the New Haven data (**Fig. S9A**) in classifying the Mannheim and the Chicago patients, respectively, into persistent and recovered. FA values corrected for age, gender, and head displacement accurately classified SBPr and SBPp patients from the Mannheim data set with an AUC = 0.67 (*p* = 0.023, tested against 10,000 random permutations, **Fig. S9B and S7D**), and patients from the Chicago data set with an AUC = 0.69 (*p* = 0.0068), (**Fig. S9C and S7E**) at baseline, and an AUC = 0.67 (p = 0.0098) (**Fig. S9D and S7F**) patients at follow-up, confirming the predictive cluster from the right SLF across sites. The application of neuroCombat significantly changes the FA values as shown in **Fig. S10** but does not change the results between groups.

### Structural Connectivity-Based Classification of SBPp and SBPr patients

We studied the structural connectivity of brain areas known to be connected by different parts of the right SLF by illustrating in 3 dimensions the white matter fiber-tracts traveling between them (**Fig.7A**). We also extracted the FA values along those tracts (**Fig.7B**). Most of the visualized white matter bundles showed thinning (i.e., decreased density) in the SBPp patients when visually inspected relative to the SBPr patients (**Fig. 7A; Fig. S11**). Additionally, there was a drop in FA along the fiber tracts in the SBPp compared to the SBPr patients with a trend towards a lower number of tracts in the former group. (**Fig.7B**).

**Figure 7.**
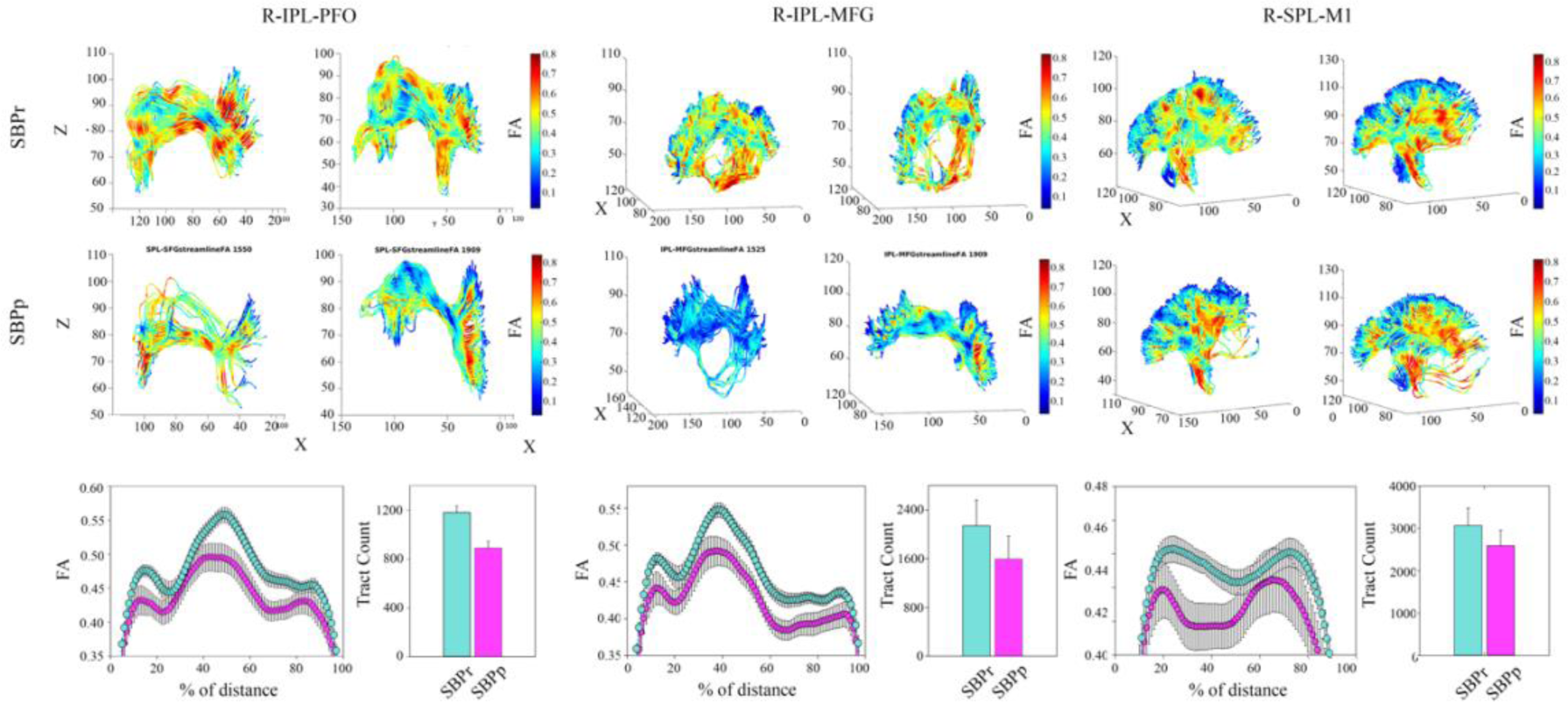
Illustration of the right SLF white matter bundles and their FA content in SBPp and SBPr patients of the New Haven data set. **(A)** Three-dimensional illustration of SLF tracts connecting the inferior parietal lobe with frontal operculum (left), inferior parietal lobe and middle frontal gyrus (middle), and superior parietal lobe and primary motor cortex (right). Upper two rows show SLF tracts connecting the two regions in 3D with FA values along those tracts shown in blue-to-red. The first row shows two representative SBPr patients and the second row shows 2 representative SBPp patients. (**B**) Plot of average FA ± SEM along the tracts depicted in A for 15 SBPr and 12 SBPp patients. The histogram plot shows the average number of tracts for these same patient groups. *Abbreviations*: IPL, inferior parietal lobe; PFO, prefrontal operculum; MFG, middle frontal gyrus; SPL, superior parietal lobe; M1, primary motor cortex.

Furthermore, using fiber count and total connected surface area connecting each pair of regions of the structural white matter connectome as the input features, we pooled the data from the three sites and machine learning cross validation to classify sub-acute back pain patients as SBPr or SBPp. The best performing structural connectivity model was based on combining 40% of the New Haven data with 100% of the Chicago data during initial training and validation and testing on the remaining 60% of the New Haven data or on the Mannheim data **(Table S7**). The average AUC (i.e., the average performance of the model) was 0.67 ± 0.03 (mean ± std) using a support vector classifier (SVC) when classifying the remainder of the New Haven sample and 0.53 ± 0.03 when classifying the Mannheim data (**Fig. S12)**.

## Discussion

The white matter properties of the right SLF were significantly different between SBPr and SBPp patients across three different sites: New Haven, Chicago, and Mannheim. Furthermore, SBPr patients showed larger FA values than pain-free controls in the right SLF in both the New Haven and Mannheim data sets. This suggests that this white matter property is a biomarker of resilience to pain chronicity where pain-free controls are composed of both high and low resilience individuals. Consistent with the concept of resilience, higher baseline fractional anisotropy in the right SLF predicted a greater percentage reduction in pain intensity at follow-up. A model based on whole-brain structural connectivity classified SBPp and SBPr patients across sites, although with a lower overall accuracy compared to the TBSS-based univariate approach. Despite the implication of the uncinate fasciculus in learning and memory,(44) which are important processes in the emergence of chronic pain,(48) we did not detect any microstructural alterations in this tract. Alterations of the uncinate fasciculus were noted in the fiber complexity(49) but not in fractional anisotropy(50) in CBP patients compared to pain-free controls. This suggests that either FA is not the appropriate measure used for studying the uncinate fasciculus in relation to pain, or that this tract is not involved in chronic aversive learning.

The SLF is a large bundle of white matter association fibers connecting occipital, temporal, and lateral parietal lobes with the ipsilateral frontal lobe.(51) In vivo studies in humans have anatomically subdivided the SLF into dorsal (SLF I), middle (SLF II), and ventral (SLF III) branches that run in parallel,(52) while the inclusion of the fourth component, the arcuate fasciculus, has been questioned by some authors due to its distinct anatomical trajectory that runs from the frontal to the temporal lobe.(53) The SLF is an essential anatomical substrate for major cognitive functions such as language (left SLF), memory, emotions, motor planning, and visuospatial processing.(54) The right SLF is particularly important in visuospatial attention to the extra-personal space and the positioning of the body in the physical space.(55) It connects frontal and parietal areas critical in the top-down control of attention during tasks involving any sensory modality.(54) The SLF cluster found in our discovery set and validated in two other independent data sets spans all three branches of the SLF white matter tract. Within the significant cluster in the discovery data set, MD was significantly increased, while RD in the right SLF was significantly decreased in SBPr compared to SBPp patients. Higher RD values, indicative of demyelination, were previously observed in chronic musculoskeletal patients across several bundles, including the superior longitudinal fasciculus.(50) Similarly, Mansour et al. found higher RD in SBPp compared to SBPr in the predictive FA cluster. While they noted decreased AD and increased MD in SBPp, suggestive of both demyelination and altered axonal tracts(42), our results show increased MD and RD in SBPr with no AD differences between SBPp and SBPr, pointing to white matter changes primarily due to myelin disruption rather than axonal loss, or more complex processes. Further studies on tissue microstructure in chronic pain development are needed to elucidate these processes.

Studies in patients with brain lesions have supported the importance of the right SLF in top-down attention processing. For example, patients who underwent glioma resection with damages to the right medial superior and middle frontal gyrus, brain regions traversed by SLF I and II, showed persistent visuospatial cognitive dysfunction postoperatively.(56) Similarly, patients with right prefrontal glioma with the resection cavity located in a region overlapping SLF I and SLF II had persistent spatial working memory deficits even in the absence of motor and language deficits.(57) Lesions within the SLF II have been linked to visuospatial neglect,(58, 59) supporting the role of this subcomponent in visuospatial awareness. Together the literature on the right SLF role in higher cognitive functions suggests, therefore, that resilience to chronic pain might be related to a top-down phenomenon involving visuospatial and body awareness.

Higher FA values in the right SLF have been associated with better performance in sustained attention in children(60) and people suffering from attention deficits or hyperactivity disorder.(61–63) After a stroke, patients who had higher baseline FA values within SLF II showed higher success rates in the visuomotor task after 4 weeks of learning compared to those with lower baseline FA values.(64) Similarly, a recent study showed that SLF II underwent plastic changes after learning of tracking movement tasks requiring both top-down attentional processes and bottom-up somatosensory feedback to adjust one’s own movement, and the degree of this plasticity predicted task-related success.(65) Since the constant adaptation of motor control in these tasks is dependent on spatial awareness and proprioception, the integrity of the SLF II appears crucial to their intact functioning. Proprioception, which is a bottom-up somatic signal closely related to visuospatial processing, is critical for recognizing one’s own body position and hence the awareness of the physical self. Changes at the higher-order level of proprioceptive processing can also affect body perception.(66) The right SLF connects a network of brain regions involved in proprioceptive awareness and allows us to perceive ourselves as separate entities from external animate and inanimate objects.(67) Lesions of this white matter bundle and/or of the right parietal lobe are associated with hemispatial neglect of the left side of extra-personal space and, in extreme cases, of one’s own body parts.(54) Specifically, SLF II and SLF III subcomponents have been linked to dysfunctions in proprioception. The inferior frontoparietal network, connected by the SLF III, showed predominant right-hemispheric activation during both the visual self-face recognition and limb proprioceptive illusion task, which suggests importance of this tract in self-awareness independent of the sensory modality and body parts involved.(68) Similarly, changes in limb position in relation to posture activate the same network, hence suggesting that constant awareness of our body position and corrective feedback on our body schema is indeed dependent on this tract.(55)

Although the findings on the direct association between impaired proprioception and CBP are inconclusive due to the variety of methods used to measure proprioceptive performance,(69–71) there is evidence that the central processing of back-related proprioceptive signals is affected in chronic low back pain.(66, 72) Indeed, it has been shown that pain perception influences body representation,(73) and body representation is altered in patients with chronic pain as investigated by psychophysical,(72, 74) and neuroimaging studies.(66, 75, 76) Interestingly, in some cases disturbances in body representation occurred before the development of chronic pain(77) and predicted decreased analgesic response to an exercise treatment,(78) suggesting a causal role of such a distortion in chronicity. While we did not collect proprioceptive and attention measures in the SBP patients, our results suggest that these measures could be useful to separate resilient and at-risk patients.

The asymmetry of the functions subserved by the SLF is notable as lesions to the right but not left SLF lead to hemispatial neglect on the left side, suggesting that vulnerabilities and strength in the neural substrate mediating attention and visuospatial processing contribute to the long-term risk or resilience to chronic pain. The observation that the structural properties of such a large association fiber network predict pain chronicity also suggests that risks and vulnerabilities to chronic pain may be determined by large, distributed frontoparietal networks on the right side of the brain. Interestingly, compared to pain-free controls, low back pain patients show functional connectivity alterations in the dorsal visual stream involved in visuospatial attention and subserved by the SLF tract, but not in the ventral visual stream.(79) The results motivated us to explore different diffusion-based parameters, primarily focusing on structural connectivity. While our SVC machine learning-based model accurately classified SBPr and SBPp patients, it did not perform better than the univariate TBSS-based analysis and was worse than the latter approach in classifying the Mannheim patients. This discrepancy is most likely because whole-brain structural connectivity approaches such as PSC(80) are more sensitive to signal-to-noise ratio and site-related differences in data acquisition parameters than TBSS such as the number of directions in the diffusion data acquisition protocol. Another explanation is that structural connectivity-based classifiers are sensitive to the heterogeneity of patients across sites because the eligibility criteria used were similar between New Haven and Chicago but different from those in Mannheim.

Our results along with previous studies suggest that resilience to chronicity of back pain may be related to the structural integrity (i.e., FA) of the right SLF. Like Mansour et al.,(42) we observed that SBPr patients show larger FA values than the pain-free controls while the SBPp patients show similar FA values to the CBP patients suggesting that the pain-free population is made up of high and low-risk groups (**Fig. 1**; **Fig. 3**). Notably, though, the DTI-based FA results reported by Mansour et al. were on the left side of the brain. While larger datasets are required to explain this discrepancy, it does suggest that resilience to chronic pain might also be a widespread brain property involving other large-scale brain networks. CBP state has been associated with white matter changes across several tracts,(50) but here we show that the SLF tract is particularly important in the transition phase of subacute to CBP state, predicting long-term chronicity in a lateralized, right-dominant manner. As pain turns chronic, it is likely to progressively involve other neural pathways, indicating changes within broader neural networks. In fact, neural signatures of sustained and chronic pain are predominately observed in somatomotor, frontoparietal, and dorsal attention networks,(34) which corresponds to the microstructural changes in tracts found in the chronic state. The right SLF, however, shows changes already at half a year into chronicity, remains stable at one year, and could therefore serve as a potential biomarker to address the need for early detection of risk.

In task-based and resting state neuroimaging studies the brain activity in the frontoparietal area was not typically observed as a risk factor for persistent pain. Instead, functional connectivity between the medial prefrontal cortex and nucleus accumbens,(39) as well as the processing of reward signals in the same pathway (40) were shown to be predictive of back pain chronicity. Investigations into structural properties showed that the risk for the development of CBP is related to a smaller nucleus accumbens and hippocampi ((39, 46, 81) in the SBPp patients relative to the other groups. Conversely, larger amygdala volume seems to be a protective factor against chronicity, as it was greater in SBPr than in both the persistent pain group and the pain-free group.(46) In the same vein, we observed greater FA values within SLF in SBPr not only compared to SBPp but also to healthy controls. The involvement of the frontoparietal area can thus be regarded as part of structural “resilience circuitry”.

Despite different time frames for the follow-up, initial (baseline) pain intensities (and accordingly different criteria for subgrouping SBP into SBPr/SBPp), population, sites, scanners, and pain questionnaires/screening used, we successfully validated results across three different sites. Moreover, we carefully addressed the potential confounding effects of head displacement, which can lead to either positive or negative bias(82) by accounting for it in the analysis. This points towards the robustness of the integrity of the SLF as a biomarker of resilience to CBP, with a potential for clinical translation.

## Limitations

Our results are based on heterogeneous samples, heterogeneous pain measures, different criteria for recovery, and different scanners across sites. In addition, at the time of analysis, we had “access” to all the data, which may lead to bias in model training and development. We believe that the data presented here is nevertheless robust since multisite validated but needs replication. Additionally, we followed standard procedures for machine learning where we never mix the training and testing sets. The models were trained on the training data with parameters selected based on cross-validation within the training data. Therefore, no models have ever seen the test data set. The model performances we reported reflect the prognostic accuracy of our model. Even though our model performance is average-to-good, which currently limits its usefulness for clinical translation, we believe that future models would further improve accuracy by using larger homogenous sample sizes and uniform acquisition sequences. Future studies could validate our results with increased sample sizes and using the same criteria across sites. In addition, our studies did not evaluate functions subserved by the right SLF such as proprioception or other types of visuo-spatial tasks. We believe that the results strongly support the future assessment of such cognitive functions in the study of risk and resilience to chronic pain.

## Conclusions and future directions

We have identified a brain white matter biomarker of resilience to CBP and validated it in multiple independent cohorts at different sites. This biomarker is easy to obtain (∼10 min of scanning time) and could open the door for translation into clinical practice, as future models on diffusion data are likely to improve accuracy by obtaining the data from larger sample sizes and using the same acquisition sequences. Although chronic pain may eventually affect other neural networks, the microstructural changes in the right SLF tract are evident early in the course of the illness and remain stable as pain progresses. Future studies should investigate how this brain structural predisposition to CBP may impact brain function, information processing, and neural networks. This could lead to potential neural targets for early interventions such as neurofeedback(83) or brain stimulation(84). In addition, cognitive and behavioral processes associated with the right SLF, such as proprioception and attentional functions, should be examined in subacute stages, as targeting these processes could add to the effective prevention of chronicity. Integrating findings from studies that used questionnaire-based tools and showed remarkable predictive power, (32) with neurobiological measures that can offer mechanistic insights into chronic pain development, could enhance predictive power in CBP prognostic modeling.

To establish the clinical usefulness of a biomarker, it should be tested in various populations, settings, and contexts, and ideally be cost-effective and simple to implement.(43) In this regard, we have introduced a promising biomarker found in brain white matter, which has considerable potential for clinical application in the prevention and treatment of CBP.

## Materials and Methods

### Data pool

#### New Haven (Discovery) data set

We recruited individuals in the New Haven, Connecticut area. Subjects were recruited through flyers and internet advertisements. All participants gave written informed consent to participate in the study. The study was approved by the Yale University Institutional Review Board.

Brain diffusion data were collected at baseline from 27 (12 females) subacute low-back pain patients (SBP, pain duration between 6-12 weeks), 29 (16 females) patients with chronic low-back pain (CLBP), and 28 (12 females) healthy controls (HC). One patient was excluded because of excessive head motion defined as >3SD from the mean Euclidian distance of either translational or rotational displacement during the MRI scanning session and one HC subject was excluded because parts of the brain were outside the field of view.

Subjects were briefly screened at first to check (1) the location of the low-back pain, (2) if they were otherwise healthy, (3) non-smokers, and (4) pain duration (between 6 and 12 weeks for SBP and more than one year for CLBP). If they passed this initial brief screen a more detailed screen was conducted where we assessed complete medical and psychiatric history. To be included in the study, SBP subjects needed to meet the criteria of having a new-onset 6 to12 weeks low back pain at an intensity more than 20/100 on the visual analogue scale (VAS) and report being pain-free in the year prior to the onset of back pain. CLBP patients had to have a pain duration of at least 1 year and a pain intensity of more than 30/100. Both SBP and CLBP participants had to (1) fulfill the International Association for the Study of Pain criteria for back pain,(85) (2) not be currently, or during the month prior to the study, on any opioid analgesics. Patients were included if their back pain was below the 12th thoracic vertebra with or without radiculopathy and was present on more days than not. SBP and CLBP diagnoses were confirmed based on history collected by an experienced clinician (P.G.). Healthy control subjects were screened likewise with, besides, the absence of any history of any pain of more than 6 weeks in duration. Participants had no history of mental disorders, chronic medical conditions (e.g., diabetes, coronary artery disease), or loss of consciousness.

The study consisted of two time points separated by approximately one year (baseline, 1-year follow-up). At each time point participants completed one testing session in the laboratory and one scanning session. Patients whose pain dropped by more than 30%(86) at follow-up were considered recovered, otherwise persistent. Of the SBP group, 12 patients were confirmed at follow-up as recovered subacute back pain patients (SBPr) and 16 as persistent subacute back pain patients (SBPp) and had diffusion tensor data collected. For demographic and clinical characteristics of the different groups see **Table S1.**

### For the MRI acquisition description **see** Supplementary Methods

#### Mannheim data set

Participants were recruited through advertisements in local newspapers and the website of the Central Institute of Mental Health in Mannheim, and patients were recruited additionally through the outpatient pain clinic of the Institute of Cognitive and Clinical Neuroscience, general practitioners, and physiotherapy practices. We determined subjects’ eligibility via a telephone screening form, which comprised questions about MRI contraindications, medication intake, current and previous drug/alcohol use, co-morbid medical and psychological conditions, and pain frequency and severity. All participants had to be between 18 and 70 years old to be eligible for study entry. Healthy controls had to be pain-free; patients with pain (SBP and CBP) had either low and/or upper back pain.

To be included in the study, SBP participants had to have a current back pain episode of 7–12 weeks duration. Patients with a current back pain episode and additional back pain episodes in their history were included as well if the episodes never exceeded 12 weeks per year. The inclusion criteria for the CBP group were a history of back pain longer than 6 months and a current back pain episode of more than 100 days. Participants were excluded if they reported any neurological disorder, psychotic episodes, current substance abuse, a major illness, contraindication for MRI, or another painful condition as the main pain problem.

Brain diffusion data were collected at baseline from 64 patients with SBP, 24 patients with CBP, and 24 healthy controls (HC). HC and patients with CBP were matched for age and gender. Two HC, three patients with CBP, and nine patients with SBP were excluded from the analysis due to excessive head motion defined as >3SD from the mean Euclidian distance of either translational or rotational displacement during the MRI scanning session. Additionally, two SBP patients’ diffusion images failed manual quality control checks due to obvious artifacts. Six patients with SBP were excluded from the analysis because they did not have interim data at follow-up. One patient with SBP was also excluded from the analysis as an outlier due to an extreme increase in pain severity from baseline to follow-up (466 percent change, M= −15,06; hence the subject was > 6 standard deviations from the sample mean). The final sample in the analysis comprised 22 HC, 21 patients with CBP, and 46 patients with SBP. Demographic and clinical characteristics of this data set, which was used for validation of our white matter biomarker, are presented in **Table S2.**

Medication use is presented in **Table S3**. All participants gave written informed consent to be involved in the study and received 10€/hour for their participation. The study was approved by the Ethics Committee of the Medical Faculty of Mannheim, Heidelberg University, and was conducted in accordance with the declaration of Helsinki in its most recent form.

#### Clinical assessments

Patients with SBP were included in the study at baseline as one group, and their pain severity was assessed at two time points (baseline, 6-month follow-up). Change in pain severity (PS) was assessed using the percentage change in the Pain Severity scale of the German version of the West Haven-Yale Multidimensional Pain Inventory(87) from baseline assessment to the follow-up screening after 6 months using the following formula: 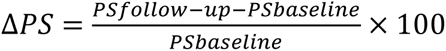. Based on the pain severity percentage change, the SBP sample was divided into recovered SBP (SBPr, *N*=28), whose pain dropped by more than 20% at follow-up and persisting SBP (SBPp, *N*=18) (Δ𝑃𝑆 > −20) following criteria already published in the literature(39) (see also **Supplementary Methods**). For the **MRI acquisition description** see **Supplementary Methods.** Comorbid mental disorders are presented in **Table S4.**

### For the MRI acquisition description **see** Supplementary Methods

#### Chicago data set (Open Pain)

The data set obtained through the OpenPain.org online database (collected in Chicago, from now on, we refer to this data set as “Chicago data set”) had 58 SBP patients (28 females) with a baseline visit (visit 1 on OpenPain, pain duration 6-12 weeks) and 60 SBP patients (29 females) with a one-year follow-up visit (visit 4 on OpenPain) on whom diffusion images were collected.

Patients were deemed recovered at one-year follow-up based on the same criterion as the New Haven data (i.e., > 30% drop in low-back pain intensity reported on the VAS). As such, we studied 35 SBPp and 23 SBPr at baseline, and 33 SBPp and 27 SBPr at follow-up. Demographic and clinical characteristics of this data set, which was used for validation of our white matter biomarker, are presented in **Table S5**. For the **MRI acquisition description** and **preprocessing of DTI Data** see **Supplementary Methods.**

### Tract-Based Spatial Statistics

Voxel-wise statistical analysis of FA was carried out using Tract-Based Spatial Statistics (TBSS)(88) part of the FSL.(89) All subjects’ FA data were then aligned into a common space (MNI standard 1-mm brain) using the nonlinear registration tool FNIRT,(90, 91) which uses a b-spline representation of the registration warp field.(92) Next, the mean FA image was created and thinned to create a mean FA skeleton, which represents the centers of all tracts common to the groups. Each subject’s aligned FA data was then projected onto this skeleton and the resulting data fed into voxel-wise cross-subject statistics. Groups (i.e., SBPr and SBPp) were compared using unpaired t-test corrected for age, gender, and motion parameters. Head displacement was estimated by eddy current correction to extract the magnitude of translations and rotations. Overall head motion was then calculated as the Euclidian distance from head translations and rotations for each subject, and these measures were Z-transformed before they were entered into the design matrix as nuisance variables. The statistical significance of TBSS-based testing was determined using a permutation-based inference(93) where the null distribution is built using 10,000 random permutations of the groups. Significance was set at *p* < 0.05, and significant clusters were identified using threshold-free cluster enhancement.(94)

### Statistical analysis

A mask formed from the significant cluster for the SBPr>SBPp contrast in the New Haven data was used to extract FA values from the Mannheim and the OpenPain data. The FA values were first corrected for confounders and then used to build a receiver operating curve (ROC) to assess classification accuracy (recovered and persistent pain patients as binary classes) based on the brain white matter data. The statistical significance of the area under the ROC (AUC) was tested against 10,000 random permutations of the group labels to generate a random distribution of the AUC values. Additionally, we tested if FA values predict pain percentage change in a dimensional approach using multiple linear regression with the FA data entered as a predictor, and pain percentage change as an outcome. In both, classification and linear regression analysis, age, gender, and motion parameters (translation and rotation) were entered as covariates of no interest.

### Estimation of structural connectivity

Structural connectivity was estimated from the diffusion tensor data using a population-based structural connectome (PSC) detailed in a previous publication.(80) PSC can utilize the geometric information of streamlines, including shape, size, and location for a better parcellation-based connectome analysis. It, therefore, preserves the geometric information, which is crucial for quantifying brain connectivity and understanding variation across subjects. We have previously shown that the PSC pipeline is robust and reproducible across large data sets.(80) PSC output uses the Desikan-Killiany atlas (DKA) (95) of cortical and sub-cortical regions of interest (ROI). The DKA parcellation comprises 68 cortical surface regions (34 nodes per hemisphere) and 19 subcortical regions. The complete list of ROIs is provided in the supplementary materials’ **Table S6**. PSC leverages a reproducible probabilistic tractography algorithm (96) to create whole-brain tractography data, integrating anatomical details from high-resolution T1 images to minimize bias in the tractography. We utilized DKA (95) to define the ROIs corresponding to the nodes in the structural connectome. For each pair of ROIs, we extracted the streamlines connecting them by following these steps: 1) dilating each gray matter ROI to include a small portion of white matter regions, 2) segmenting streamlines connecting multiple ROIs to extract the correct and complete pathway, and 3) removing apparent outlier streamlines. Due to its widespread use in brain imaging studies(97, 98), we examined the mean fractional anisotropy (FA) value along streamlines and the count of streamlines in this work. The output we used includes fiber count, fiber length, and fiber volume shared between the ROIs in addition to measures of fractional anisotropy and mean diffusivity.

### Machine learning and cross-validation

The processing steps applied to the PSC output are summarized in **Fig. S1** and were published previously.(80) We used fiber count and total connected surface area connecting each pair of regions as the input features, which were 87 x 87 connectivity matrices of the DKA ROIs; the latter two features were chosen because of their demonstrated reliability in prior work.(80) Dimensions were reduced using tensor principal component analysis (TNPCA) with hyperparameter K = 87. Next, because data originated from different sites (New Haven, Chicago, and Mannheim), we applied the ComBat harmonization(47) to account for the effect of different sites on the data. Linear regression was used next to correct the data for age, gender, and head translation and rotation magnitudes. We used logistic regression, linear support vector classification (SVC), and random forest as our regression methods with leave-one-out cross-validation (LOOCV) and the number of components as a hyperparameter. Hence, the number of PCs included in the model was randomly chosen from a sequence of 22 numbers starting with 1 with an increment of 4 and a maximum of 87. The training of our regression models was performed using different combinations of data originating from New Haven and the Chicago database as outlined in **Table S7** and tested on the remainder of the data from these two sites or on the data originating from Mannheim. The combination of the data during model training was bootstrapped 50 times and testing and validation were repeated accordingly. We chose to train the model only on the data originating from New Haven and Chicago because the eligibility criteria were quite similar between the two sets but different from those used to recruit SBP patients at Mannheim. The average performance of our classifier was based on the models that achieved a validation area under (AUC) the receiver operating characteristic curve (ROC) of ≥ 0.75 during training. Of note, such models cannot tell us the features that are important in classifying the groups. Hence, our model is considered a black-box predictive model like neural networks.

## Acknowledgments and funding sources

The New Haven study was funded by the National Institute on Drug Abuse grant number K08DA037525. The Mannheim study was supported by a grant from the Deutsche Forschungsgemeinschaft (SFB1158/B03 to F.N. and H.F.) and it has been registered on the ‘‘German Clinical Trials Register’’ the registration ID is DRKS00008835. The access website is as follows: https://www.drks.de/. The OPP project (Principal Investigator: A. Vania Apkarian, Ph.D. at Northwestern University) is supported by the National Institute of Neurological Disorders and Stroke (NINDS) and National Institute on Drug Abuse (NIDA).

## Data availability

New Haven and Mannheim data sets are available upon reasonable request from the lead contacts (<u>)</u>, as the studies are ongoing. The imaging data set, demographical, and all clinical variables will be shared in an anonymized form to ensure participant privacy and to comply with legal requirements. The data set obtained from Chicago is from the OpenPain Project (OPP) publicly available at: https://www.openpain.org/. Connectivity analysis scripts were written in R (version 4.3.1) and are shared on https://github.com/MISICMINA/DTI-Study-Resilience-to-CBP.git.

## Author Contributions

Study concept and design, H.F., F.N., P.G., and M.M.; acquisition of data, M.M., K.U., and M.L.; analysis and interpretation of data, M.M. and N.L.; drafting of the manuscript, M.M.; critical revision of the manuscript for important intellectual content, M.M., N.L., F.Z., K.S., M.L., K.U., M.N.U., A.F., G.S., Z.Z., F.N., P.G., and H.F.; obtained funding, P.G., H.F., and F.N.; administrative or technical support, M.M., F.Z., study supervision, F.N., P.G., and H.F.

