## Supplementary Information for "Brain white matter pathways of resilience to chronic back pain: a multisite validation"

**This PDF file includes:**

Supplementary Methods
Figures S1 to S12
Tables S1 to S7
SI References

### Supplementary Methods

#### *New Haven data set*

**Magnetic resonance imaging.** Participants underwent an anatomical T1-weighted scan and two consecutive 2.5-minute-long diffusion tensor imaging (DTI) scans in the same session. A Siemens 3 T Trio B magnet (Erlangen, Germany) equipped with a 32-channel head coil was used to acquire the images. The 3D magnetization prepared rapid gradient echo (MPRAGE) T1-weighted acquisition sequence was as follows: repetition time/echo time (TR/TE) = 1900/2.52 ms, flip angle = 9°, matrix size = 256 x 256, number of slices = 176, image resolution = = $1 \times 1 \times 1$ mm$^{3.}$ The diffusion-weighted images were acquired using spin-echo echo planar imaging (SE-EPI) sequence using the following scan parameters: TR/TE = 2200/84.0 ms, flip angle = 90°, matrix size = $110 \times 110 \times 64$, multi-band acceleration factor = 4, image resolution = $2 \times 2 \times 2$ mm$^3$. Diffusion gradients were applied along 64 directions with a b-value of 1000 s/mm$^2$. For each set of DTI data, one volume with no diffusion weighing (i.e., b=0 s/mm$^2$) was acquired at the beginning of the scan.

#### *Mannheim data set*

**Participants.** A trained psychologist interviewed all participants to assess comorbid mental disorders using the German version of the Structured Clinical Interviews (SCID I) for the Diagnostic and Statistical Manual of Mental Disorders (DSM IV)(1) **Supplementary Table 2**). In addition, all subjects completed the German versions of the Chronic Pain Grade (CPG),(2) the Örebro Musculoskeletal Pain Questionnaire (OMPQ yellow flag),(3) the Hospital Anxiety and Depression Scale (HADS),(4) and the Perceived Stress Scale (PSS).(5)

**Magnetic resonance imaging.** A 3 Tesla Tim TRIO whole body scanner (SIEMENS Healthineers, Erlangen, Germany), equipped with a 12-channel head coil was used to acquire the images. Shimming of the scanner was done to account for maximum magnetic field homogeneity. Participants underwent an anatomical T1-weighted MPRAGE imaging scan with the following sequence: TR/TE = 2300/ 2.98 ms, flip angle = 9°, matrix size = $240 \times 256$, number of slices = 192, image resolution = $1 \times 1 \times 1$ mm3. The diffusion weighted images were acquired using SE-EPI sequence with the scan parameters as follows: TR/TE = 7400/85 ms, matrix size = $220 \times 256 \times 30$, GRAPPA = 2, image resolution = $2 \times 2 \times 2$ mm$^3$. Diffusion gradients were applied along 30 directions using a b-value of 1000 s/mm2. One volume with no diffusion weighing was acquired at the beginning of the scan.

#### *Open Pain (Chicago) data set*

**Magnetic resonance imaging.** Part of this data has been already published by Mansour et al.(6) Therefore, we will briefly describe acquisition parameters. MPRAGE type T1-anatomical brain images were acquired with a 3T Siemens Trio whole-body scanner with echo-planar imaging capability using the standard radio-frequency head coil with the following parameters: TR/TE = 2,500/3.36ms, flip angle = 9°, matrix size = 256 × 256; number of slices = 160, image resolution = 1 × 1 ×1 $mm^3$. DTI images were acquired on the same day using SE-EPI with TR/TE =9000/83, flip angle = 90 degrees, in-plane matrix resolution = 112 × 130; 73 slices; image resolution = 2 × 2 × 2 $mm^3$. Images had an isotropic distribution along 60 directions using a b value of 1000 s/$mm^2$. For each set of diffusion-tensor data, 8 volumes with no diffusion weighting were acquired at equidistant points throughout the acquisition.

**Preprocessing of DTI Data.** Preprocessing of all data sets was performed employing the same procedures and the FMRIB diffusion toolbox (FDT) running on FSL version 6.0. (7) First, diffusion-weighted data were visually inspected for obvious artifacts or missing parts of the brain. Next, the data were corrected for eddy currents and head motion by employing affine registration to the no diffusion volume using eddy_openmp from the FSL toolbox. *Eddy current* corrects for image distortions due to susceptibility-induced distortions and eddy currents in the gradient coils.(8) We do note, however, that as we did not acquire data in the phase-opposite direction, the susceptibility-induced distortions may not be fully corrected. Brain images were then skull stripped and a diffusion tensor model was fit, using FMRIB Diffusion Toolbox (FDT) part of FSL(9), at each voxel to calculate the fractional anisotropy.(10, 11) FA reflects the degree of water diffusion within a voxel with values ranging between 0 and 1 where large values indicate directional dependence of Brownian motion due to white matter tracts and small values indicate more isotropic diffusion and less directionality.(12)

**Harmonization of DTI data using neuroCombat**. Because the 3 data sets originated from different sites using different MR data acquisition parameters and slightly different recruitment criteria, we applied neuroCombat (13) to correct for site effects and then repeated the TBSS analysis shown in Figure 1 and the validation analyses shown in Figures 5 and 6. First, the FA maps derived using FDT toolbox were pooled into one TBSS analysis where registration to a standard template FA template (FMRIB58_FA_1mm.nii.gz part of FSL) was performed. Next, neuroCombat was applied to the FA maps as implemented in Python with batch (i.e., site) effect modeled with a vector containing 1 for New Haven, 2 for Chicago, and 3 for Mannheim originating maps respectively. The harmonized maps were then skeletonized to allow for TBSS.

**Supplementary Figures**

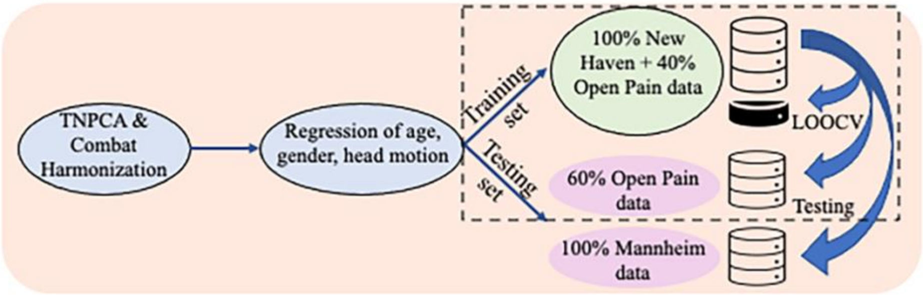


**Fig. S1.** Schematic representation of the sequential steps from left to right performed in structural connectivity data preparation, correction for confounders, and machine learning model building and testing. The dashed rectangle indicates that the combination of the data during model training was bootstrapped 50 times and validation and testing were repeated accordingly.

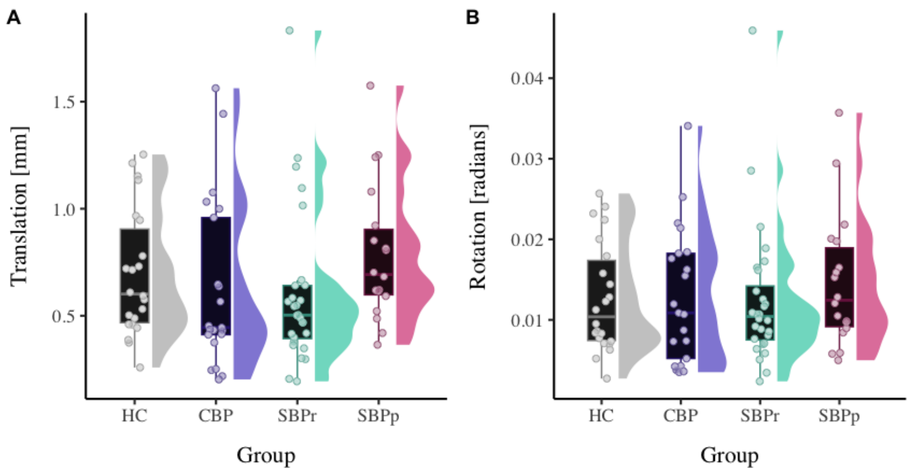


**Fig. S2.** Head displacement (translation (**A**) and rotation (**B**) parameters) estimated by eddy current correction per each group in the Discovery data set.

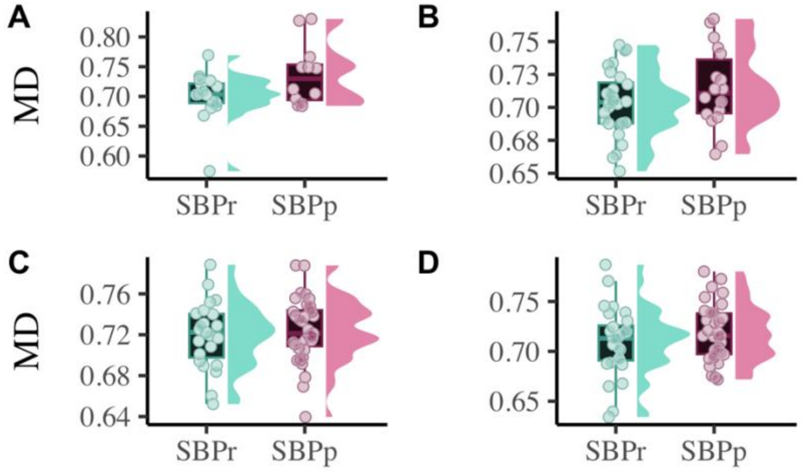


**Fig. S3.** Distribution plots of mean diffusivity (MD) extracted from the RSLF mask shown in **Fig 1**. The right SLF MD is significantly increased ($p < 0.05$) in the SBPr compared to SBPp patients in the New Haven data (**A**). The RSLF MD is not significantly different for the Mannheim data ($p = 0.12$) (**B**) nor for the Chicago data neither on visit 1 ($p = 0.44$) **(C)** nor on visit 4 ($p = 0.38$) **(D)**. The MD values were compared between groups after accounting for age, sex, and motion parameters. The values on the plot are scaled up by $10^{-4}$ for graphical depiction.

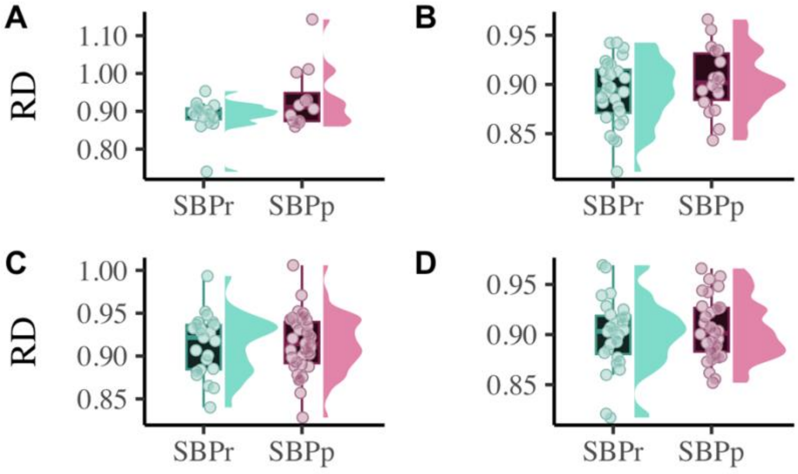


**Fig. S4.** Distribution plots of radial diffusivity (RD) extracted from the RSLF mask shown in **Fig 1**. The right SLF RD is significantly decreased ($p < 0.05$) in the SBPr compared to SBPp patients in the New Haven data (**A**). The RSLF RD is not significantly different for the Mannheim data ($p = 0.24$) (**B**). The RSLF RD is not significantly different in the Chicago data neither on visit 1 ($p = 0.78$) (**C)** nor on visit 4 ($p = 0.70$) (**D**). The RD values were compared between groups after accounting for age, sex, and motion parameters. The values on the plot are scaled up by $10^{-4}$ for graphical depiction.

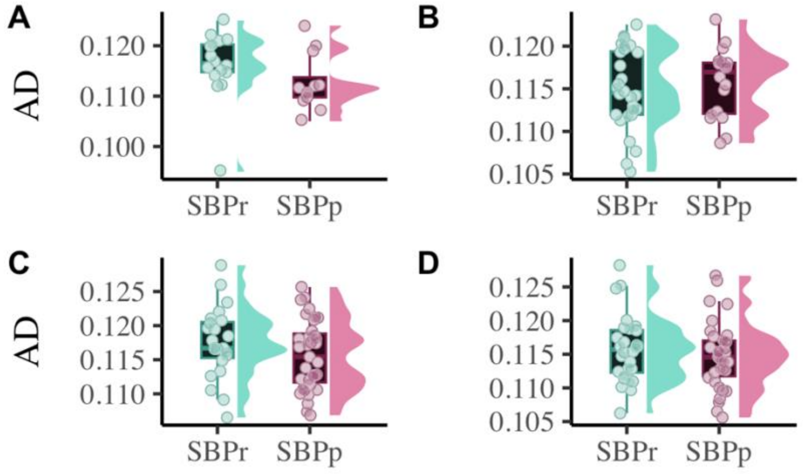


**Fig. S5.** Distribution plots of axial diffusivity (AD) extracted from the RSLF mask shown in **Fig 1**. The RSLF AD is not significantly different (p = 0.28) in the SBPr compared to SBPp patients in the New Haven data (**A**). The RSLF AD (p = 0.62) is not significantly different for the Mannheim data (**B**) nor for the Chicago data neither on visit 1 (p = 0.15) (**C**) nor on visit 4 (p = 0.36) (**D**). The AD values were compared between groups after accounting for age, sex, and motion parameters. The values on the plot are scaled up by $10^{-3}$ for graphical depiction.

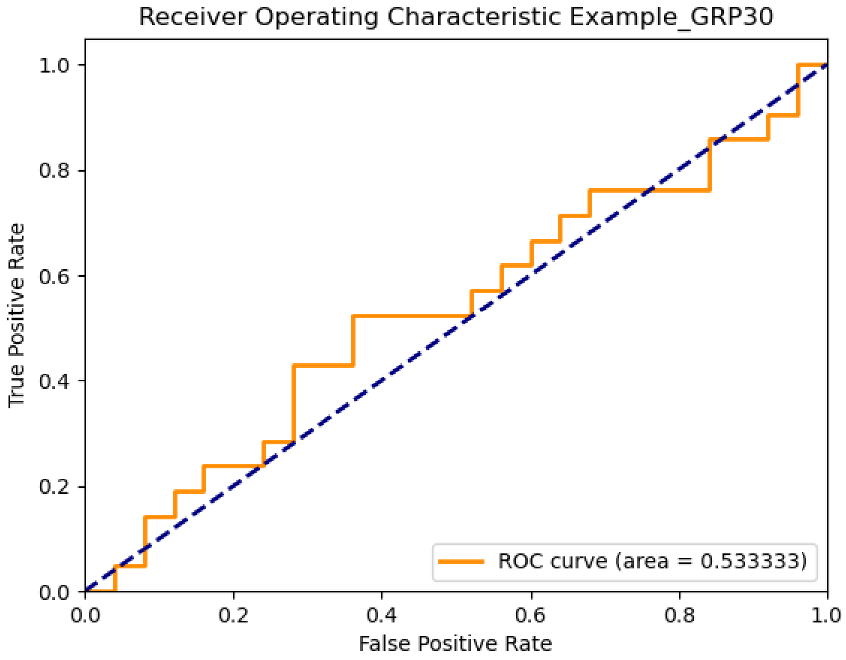


**Fig. S6.** The receiver operating characteristic curve shows the area under the curve (AUC=0.53) when using the mask from the New Haven data to classify the SBPr and SBPp patients from Mannheim using the 30% recovery criterion.

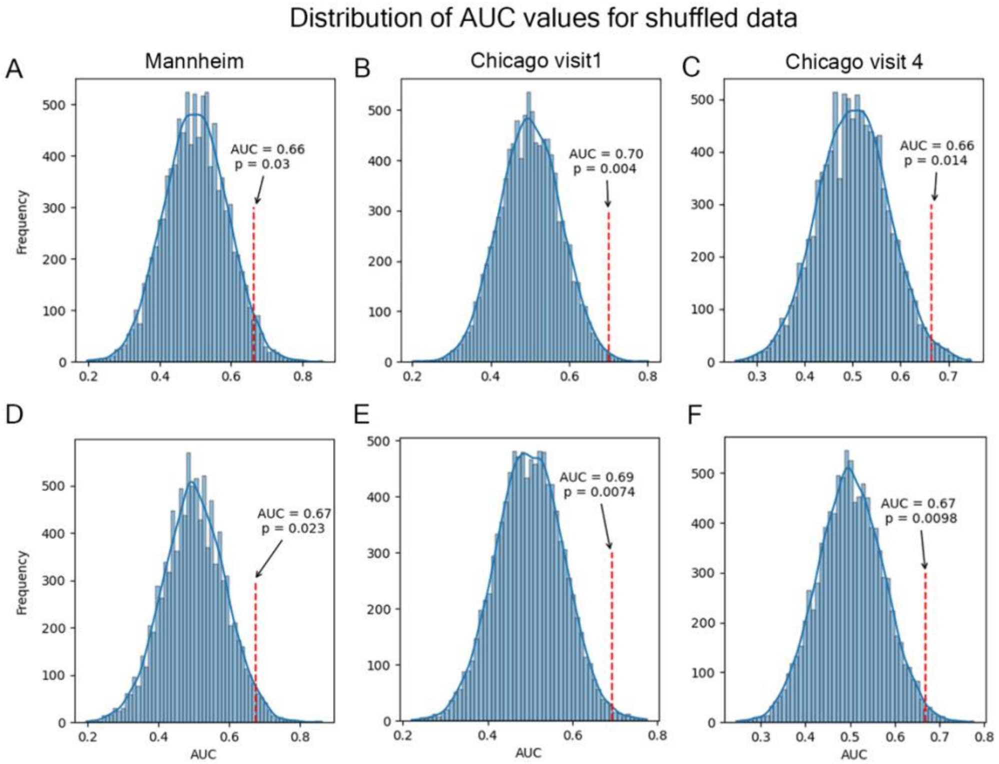


**Fig. S7.** Distribution of the values of the area under the curve (AUC) when shuffling the label of the groups (recovered, persistent) using the FA values from the right SLF before (**A-C**) and after (**D-F**) neuroCombat harmonization. The distributions for the Mannheim data are shown in **A** and **D**; the distributions of the Chicago data sets are shown in **B** and **E** and **C** and **F** respectively. The labels were shuffled 10,000 times; the p-values represent the probability of obtaining an AUC larger than the one obtained from the non-shuffled labels.

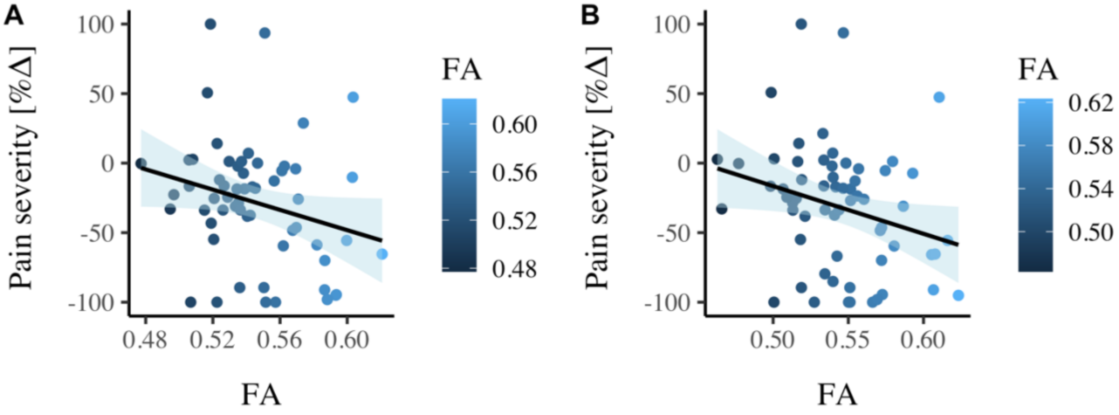


**Fig. S8**. Correlation between FA values and pain severity with higher FA values (greater structural integrity) associated with the greater reduction in pain (percentage change) at baseline (**A**) and follow-up (**B**) in the Chicago data.

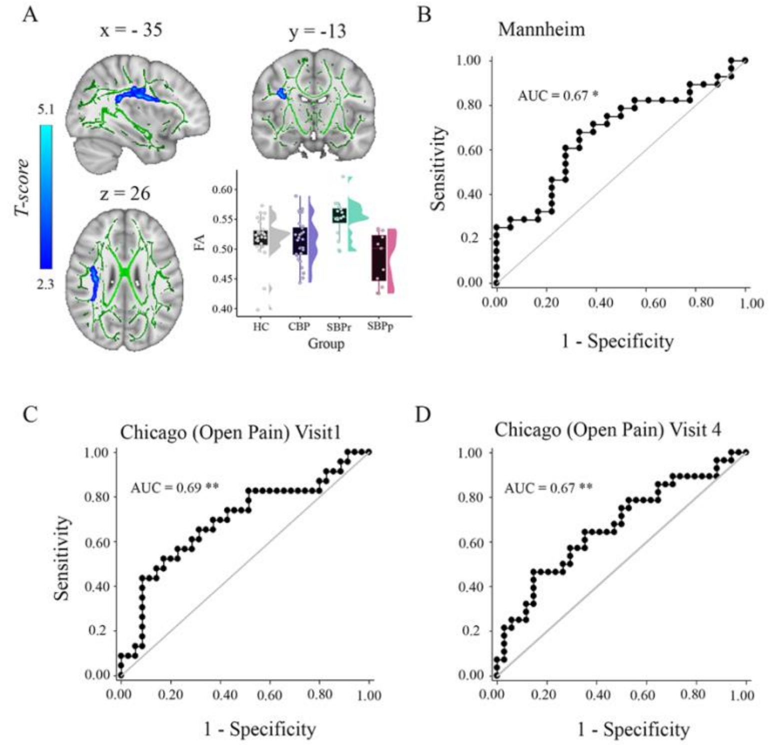


**Fig. S9**. TBSS analysis and RSLF FA validation after applying neuroCombat after pooling subjects from the 3 sites into one analysis. (**A**) A whole brain comparison over the white matter skeleton between SBPr and SBPp patients at baseline and distribution of FA values for each group (New Haven data set). Results of unpaired t-test ($p < 0.05$, 10,000 permutations) showing significantly increased FA (fractional anisotropy) in SBPr (recovered) compared to SBPp (persistent) patients within the right superior longitudinal fasciculus (SLF) in the New Haven data set. (**B-D**) ROC shows the AUC when using the mask in A to classify the SBPr and SBPp patients at each of the respective sites. *, $p < 0.05$; **, $p < 0.01$.

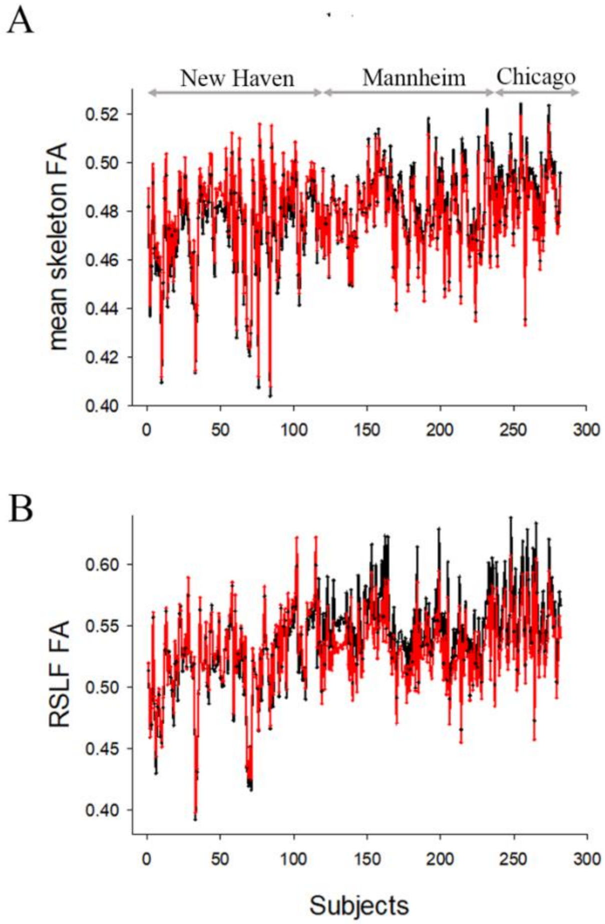


**Fig. S10**. Plot of FA values before (black) and after (red) applying harmonization using neuroCombat calculated as mean skeletal FA (**A**) and right SLF FA (**B**). Each subject's (x-axis) FA value (y-axis) is plotted for the 3 data sets Chicago, New Haven, and Manheim from left to right. FA values significantly changed when we compared the mean skeletal FA or right SLF FA before to after neuroCombat (paired t-test, $p < 10^{-7}$).

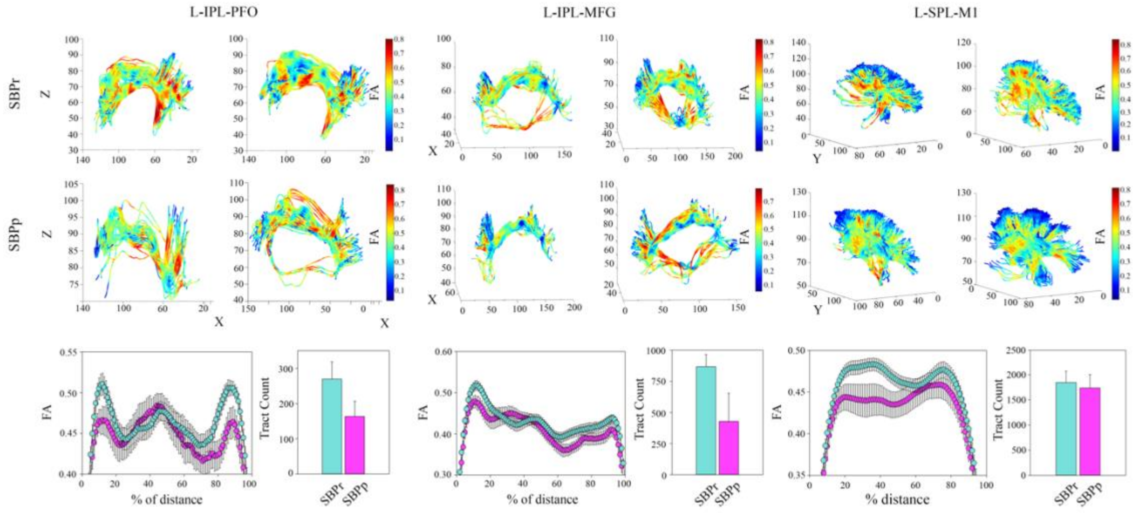


**Fig. S11. (A)** Three-dimensional illustration of the left (L) SLF tracts connecting the inferior parietal lobe with prefrontal operculum (left), inferior parietal lobe with middle frontal gyrus (middle), and superior parietal lobe and primary motor cortex (right). Upper two rows show SLF tracts connecting the two regions in 3-D with FA values along those tracts shown in blue-to-red. The first row shows two representative SBPr patients and the second row shows 2 representative SBPp patients. (**B**) Plot of average FA ± SEM along the tracts depicted in **A** for 15 SBPr and 12 SBPp patients. The histogram plot shows the average number of tracts for these same patient groups. *Abbreviations*: IPL, inferior parietal lobe; PFO, prefrontal operculum; MFG, middle frontal gyrus; SPL, superior parietal lobe; M1, primary motor cortex.

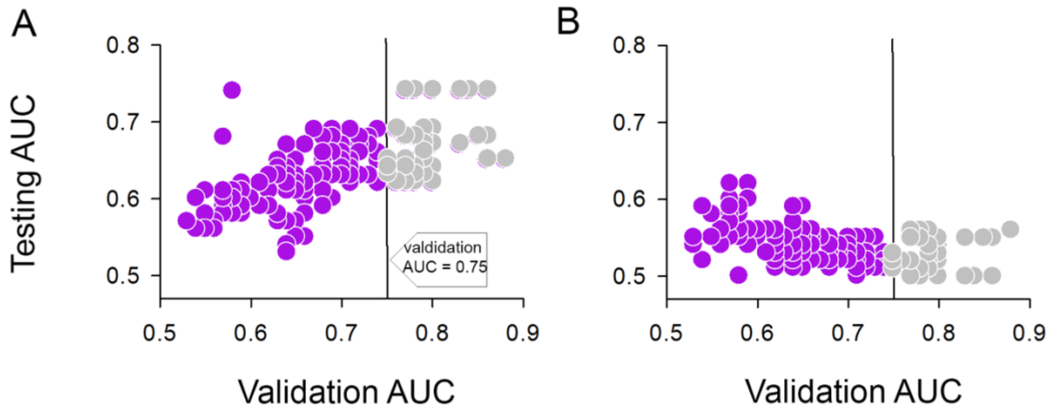


**Fig. S12.** Scatterplot of the testing AUC vs. the model validation AUC for all the machine learning models applied to the structural connectivity data. A combination of 40% of the New Haven data and 100% of the Open Pain data was used for model training and validation Testing was performed on 60% of the New Haven data (**A**), and the Mannheim data (**B**) Each point is the result of averaging the performance of the models (i.e., AUC) across 50 iterations as the combination of the data in model training stage was bootstrapped 50 times. The green circles were the final models considered in the result sections with an initial validation AUC ≥ 0.75.

### Supplementary Tables

**Table S1. New Haven sample characteristics**

| | **HC** (*N*=28) | **CLBP** (*N*=28) | **SBPr** (*N*=16) | **SBPp** (*N*=12) | **t(df)‡, p-value** | **Missing** |
|---|---|---|---|---|---|---|
| **Age (years)** | 30.1(10.1) | 30.7 (11.9) | 30.8 (8.8) | 38.0 (12.5) | + 1.80(26), 0.08 | 0/0/0/0 |
| **Gender (m/f)** | 16/12 | 12/16 | 11/5 | 7/5 | + 0.32(1), 0.57† | 0/0/0/0 |
| **BMI (Kg/m2)** | 24.1 (3.6) | 24.2 (3.7) | 25.5 (5.2) | 26.2 (4.5) | + 0.35(26), 0.73 | 0/0/0/0 |
| **Delta pain severity: absolute** | NA | NA | - 25.4 (15.4) | 8.0 (17.2) | **+ 5.0(26), $<10^{-4}$*** | NA/NA/0/0 |
| **Delta pain severity: percentage** | NA | NA | - 66.3 (26.9) | 39.1 (68.1) | **+ 5.7(26), $<10^{-5}$*** | NA/NA/0/0 |
| **Pain Duration** | NA | 5.3 (4.7) | 8.6 (3.6) | 10.9 (3.1) | + 1.87(26), 0.08 | 0/0/0/0 |
| **Pain Intensity** | NA | 4.5 (2.0) | 36.7 (18.8) | 33.7 (15.9) | - 0.45(26), 0.66 | NA/0/0/0 |
| **BDI** | 2.6 (3.3) | 6.4 (6.0) | 7.4 (4.8) | 3.1 (3.8) | **- 2.65(26), 0.02*** | 0/0/0/0 |
| **BAI** | 3.4 (5.7) | 7.6 (8.2) | 6.4 (6.8) | 4.8 (2.8) | - 0.75(26), 0.46 | 0/0/0/0 |
| **MPQ** | NA | 10.6 (4.6) | 9.2 (5.2) | 9.1 (4.2) | - 0.08(26), 0.93 | NA/0/0/0 |
| **PCS** | NA | 15.2 (9.8) | 12.8 (11.4) | 11.3 (8.1) | - 0.40(25),0.69 | NA/0/1/0 |

‡, t-score (degrees of freedom); †, Chi-square test; *, $p < 0.05$; Abbreviations: BMI, body mass index; BDI, Beck's Depression Index; BAI, Beck's Anxiety Index; MPQ, McGill Pain Questionnaire; PCS, Pain Catastrophizing Scale. Values show the group mean and standard deviation in parenthesis.

**Table S2. Mannheim sample characteristics**

| | HC (*N*=22) | CBP (*N*=21) | SBPr (*N*=28) | SBPp (*N*=18) | t(df)‡, p-value | Missing |
|---|---|---|---|---|---|---|
| **Age in years** | 36.9 (14.4) | 40.0 (16.0) | 32.8 (12.0) | 32.1 (11.2) | +0.19(38.2), 0.85 | 0/0/0/0 |
| **Gender (m/f)** | 12/10 | 11/10 | 8/20 | 6/12 | +0.0002(1),0.99‡ | 0/0/0/0 |
| **Number of days with pain during last year** | NA | 247 (84.0) | 74.8 (43.2) | 72.4 (34.9) | +0.2(41.5,2), 0.84 | NA/1/0/0 |
| **Delta pain severity (FU-BL): absolute** | NA | NA | -1.90 (1.18) | 0.19 (0.71) | -7.47(43.8), $<10^{-9}$* | NA/NA/0/0 |
| **Delta pain severity (FU-BL): percentage** | NA | NA | -50.9 (27.3) | 8.73 (26.0) | -7.45(37.7), $<10^{-9}$* | NA/NA/0/0 |
| **Pain severity (MPI)** | NA | 4.92 (1.38) | 3.80 (1.43) | 3.78 (1.54) | +0.06(34.5), 0.95 | NA/1/0/0 |
| **Interference (MPI)** | NA | 2.43 (1.23) | 1.48 (1.01) | 1.49 (1.00) | -0.03(32), 0.97 | 3/0/2/2 |
| **Negative mood (MPI)** | NA | 2.83 (1.07) | 2.40 (1.08) | 2.71 (1.15) | -0.87(30.5), 0.39 | 3/0/2/2 |
| **Life control (MPI)** | NA | 3.97 (0.971) | 4.05 (1.24) | 3.54 (1.23) | +1.3(32), 0.2 | 3/0/2/2 |
| **Support (MPI)** | NA | 2.60 (1.85) | 1.85 (1.28) | 1.56 (1.41) | +0.66(29.4), 0.52 | 3/0/2/2 |
| **ÖMPQ** | NA | 78.1 (19.5) | 63.6 (21.1) | 69.7 (16.1) | -1.06(37.9), 0.3) | 12/1/2/2 |
| **CPG**[a] | NA | 1.00 [0, 6.00] | 0 [0, 4.00] | 0 [0, 6.00] | -0.65(25.9), 0.52 | 9/0/0/2 |
| **Active coping (PRSS)** | NA | 3.30 (0.74) | 2.94 (0.96) | 3.14 (0.66) | -0.76(36.6), 0.45 | NA/6/4/3 |
| **Catastrophizing (PRSS)** | NA | 1.35 (0.87) | 1.09 (0.73) | 1.10 (0.76) | -0.02(28.9), 0.98 | NA/6/4/3 |
| **Anxiety (HADS)** | 4.37 (2.39) | 8.24 (4.47) | 7.27 (4.63) | 7.47 (3.60) | -0.15(35.3), 0.88 | 3/0/2/3 |
| **Depression (HADS)** | 6.21 (6.12) | 6.00 (4.40) | 4.42 (4.23) | 4.60 (2.85) | -0.15(37.9), 0.87 | 3/0/2/3 |
| **Perceived stress (PSS)** | 4.74 (5.00) | 11.0 (5.23) | 12.0 (5.25) | 11.4 (5.07) | +0.34(32.7), 0.73 | 3/0/2/2 |

‡, t-score (degrees of freedom); †, Chi-square test; *, $p < 0.05$; MPI, West Haven-Yale Multidimensional Pain Inventory; CPG, Chronic Pain Grade; ÖMPQ, Örebro Musculoskeletal Pain Questionnaire; PRSS, pain related self-statements; HADS, Hospital Anxiety and Depression Scale; HC, healthy control; CBP, chronic back pain; SBP, patients with subacute back pain; SBPp/r, patients with SBP with persistent pain or recovered pain after 6 months; FU, follow-up after 6 months; BL, baseline assessment; SD, standard deviation. Values show the group mean and standard deviation in parenthesis. [a]For the Chronic Pain Grade the median and interquartile range are shown.

**Table S3. Reported medication use (Mannheim sample)**

| | Regular use | Occasional use | Past use |
|---|---|---|---|
| NSAIDs | 6 SBP,3 CBP | 1 HC, 2 SBP, 3 CBP | 1 HC, 22 SBP, 14 CBP |
| Statins | 2 CBP | / | / |
| Angiotensin receptor blockers | 2 SBP | 1 CBP | / |
| Proton-pump inhibitors | 1 SBP | / | / |
| Antihistamines | 1 CBP | 1 HC, 1 CBP | / |
| Benzodiazepines | / | 1 CBP | 1 SBP, 2 CBP |
| ACE inhibitors | / | / | 1 CBP |
| Cannabinoids | / | / | 1 SBP |

**Table S4. Comorbid mental disorders (Mannheim sample); diagnoses according to the Diagnostic and Statistical Manual of Mental Disorders IV (DSM IV)**

| | Code | Diagnosis | Acute | Remitted |
|---|---|---|---|---|
| SBP | 296.26 | Major depressive disorder, single episode | | 10 |
| | 296.30/296.36 | Major depressive disorder, recurrent | 1 | 1 |
| | 300.29 | Specific phobia | 1 | |
| | 300.30 | Obsessive-compulsive disorder | | 1 |
| | 305.xx | Abuse: Opioids/Amphetamine/Cannabis/ Sedative-, hypnotic-, or anxiolytic-related | | 1 |
| | 307.10 | Anorexia Nervosa | | 2 |
| | 307.51 | Bulimia Nervosa | | 2 |
| | 309.81 | Posttraumatic stress disorder | | 2 |
| CBP | 296.26 | Major depressive disorder, single episode | | 2 |
| | 296.33 | Major depressive disorder, recurrent severe without psychotic features | | 2 |
| | 296.36 | Major depressive disorder, recurrent | 2 | 1 |
| | 300.01 | Panic disorder, without agoraphobia | 2 | |
| | 300.22 | Agoraphobia without history of panic disorder | | 1 |
| | 303.90 | Dependence: Alcohol | | 1 |
| | 304.10 | Dependence: Sedative-, hypnotic-, or anxiolytic-related | | 1 |
| | 304.30 | Dependence: Cannabis | | 1 |
| | 305.xx | Abuse: Opioids/Amphetamine/Cannabis/ Sedative-, hypnotic-, or anxiolytic-related | | 1 |
| | 307.10 | Anorexia Nervosa | 1 | |
| | 307.51 | Bulimia Nervosa | | 1 |

**Table S5. Chicago (Open Pain) sample characteristics**

| | SBPr (*N*=23) | SBPp (*N*=35) | t(df) ‡, p-value | SBPr (*N*=28) | SBPp (*N*=34) | t(df)‡, p-value | Missing |
|---|---|---|---|---|---|---|---|
| **Age (years)** | 41.7 (12.0) | 43.6 (9.3) | +0.7(56), 0.48 | 43.7 (11.5) | 45.3 (9.6) | + 0.61(60),0.54 | 0/0/0/0 |
| **Gender (m/f)** | 12/9 | 16/19 | 1.6(1),0.2† | 15/13 | 17/17 | 0.08(1), 0.78† | 0/0/0/0 |
| **Delta pain severity: absolute** | - 40.6 (20.8) | -3.4 (15.6) | + 7.8(56), $<10^{-6}$ * | - 43.1 (20.6) | - 3.2(15.6) | **+ 8.7(60), $<10^{-6}$*** | 0/0/0/0 |
| **Delta pain severity: percentage** | - 68.9 (26.2) | -1.6 (31.9) | + 8.4(56), $<10^{-6}$ * | - 69.7 (25.9) | 0.3 (31.6) | **+ 9.3(60), $< 10^{-6}$*** | 0/0/0/0 |
| **Pain Duration (weeks)** | 9.9 (4.1) | 8.4 (4.3) | - 1.3(55), 0.18 | 67.8 (5.6) | 65.2 (5.7) | - 1.8(60), 0.07 | 0/1/0/0 |
| **Pain Intensity** | 58.0 (15.2) | 67.7 (17.2) | + 2.3(56),0.03* | 18.0 (16.8) | 65.7(15.8) | **+ 11.5(60), $<10^{-6}$*** | 0/0/0/0 |
| **BDI** | 5.7 (5.2) | 7.3 (4.6) | + 1.0(39), 0.31 | 6.3 (6.6) | 16.0 (9.5) | + 1.4(42), 0.16 | 7/10/10/8 |
| **MPQ** | 10.9 (4.5) | 18.2 (17.9) | + 1.9(54), 0.06 | 13.4 (26.3) | 16.0 (9.5) | **+ 4.3 (56), $<10^{-4}$ *** | 0/2/3/1 |

‡, t-score (degrees of freedom); †, Chi-square test; *, $p < 0.05$; See Tables 1& 2 for abbreviations.

**Table S6. The complete list of the cortical and subcortical regions based on the Desikan-Killiany atlas (DKA) used to define the regions of interest (ROI) corresponding to the nodes in the structural connectivity analysis.**

| Subcortical regions | Cortical regions |
|---|---|
| LH_Lateral-Ventricle | ctx-lh-bankssts |
| LH_Cerebellum-Cortex | ctx-lh-caudalanteriorcingulate |
| LH_Thalamus-Proper | ctx-lh-caudalmiddlefrontal |
| LH_Caudate | ctx-lh-cuneus |
| LH_Putamen | ctx-lh-entorhinal |
| LH_Pallidum | ctx-lh-fusiform |
| LH_Hippocampus | ctx-lh-inferiorparietal |
| LH_Amygdala | ctx-lh-inferiortemporal |
| LH_Accumbens | ctx-lh-isthmuscingulate |
| RH_Lateral-Ventricle | ctx-lh-lateraloccipital |
| RH_Cerebellum-Cortex | ctx-lh-lateralorbitofrontal |
| RH_Thalamus-Proper | ctx-lh-lingual |
| RH_Caudate | ctx-lh-medialorbitofrontal |
| RH_Putamen | ctx-lh-middletemporal |
| RH_Pallidum | ctx-lh-parahippocampal |
| RH_Hippocampus | ctx-lh-paracentral |
| RH_Amygdala | ctx-lh-parsopercularis |
| RH_Accumbens | ctx-lh-parsorbitalis |
| | ctx-lh-parstriangularis |
| | ctx-lh-pericalcarine |
| | ctx-lh-postcentral |
| | ctx-lh-posteriorcingulate |
| | ctx-lh-precentral |
| | ctx-lh-precuneus |
| | ctx-lh-rostralanteriorcingulate |
| | ctx-lh-rostralmiddlefrontal |
| | ctx-lh-superiorfrontal |
| | ctx-lh-superiorparietal |
| | ctx-lh-superiortemporal |
| | ctx-lh-supramarginal |
| | ctx-lh-frontalpole |
| | ctx-lh-temporalpole |
| | ctx-lh-transversetemporal |
| | ctx-lh-insula |
| | ctx-rh-bankssts |
| | ctx-rh-caudalanteriorcingulate |
| | ctx-rh-caudalmiddlefrontal |
| | ctx-rh-cuneus |
| | ctx-rh-entorhinal |

| | |
|---|---|
| | ctx-rh-fusiform |
| | ctx-rh-inferiorparietal |
| | ctx-rh-inferiortemporal |
| | ctx-rh-isthmuscingulate |
| | ctx-rh-lateraloccipital |
| | ctx-rh-lateralorbitofrontal |
| | ctx-rh-lingual |
| | ctx-rh-medialorbitofrontal |
| | ctx-rh-middletemporal |
| | ctx-rh-parahippocampal |
| | ctx-rh-paracentral |
| | ctx-rh-parsopercularis |
| | ctx-rh-parsorbitalis |
| | ctx-rh-parstriangularis |
| | ctx-rh-pericalcarine |
| | ctx-rh-postcentral |
| | ctx-rh-posteriorcingulate |
| | ctx-rh-precentral |
| | ctx-rh-precuneus |
| | ctx-rh-rostralanteriorcingulate |
| | ctx-rh-rostralmiddlefrontal |
| | ctx-rh-superiorfrontal |
| | ctx-rh-superiorparietal |
| | ctx-rh-superiortemporal |
| | ctx-rh-supramarginal |
| | ctx-rh-frontalpole |
| | ctx-rh-temporalpole |
| | ctx-rh-transversetemporal |
| | ctx-rh-insula |

**Table S7. Summary of the type of data combinations expressed in % of subjects from each site used to build and test the brain connectivity-based machine learning models to classify recovered and persistent SBP patients.**

| **Training & Validation**[‡] | | **Testing**[‡] | | |
|---|---|---|---|---|
| *New Haven* | *Chicago* | *New Haven* | *Chicago* | *Mannheim* |
| 100% | 0% | 0% | 100% | 100% |
| 100% | 40% | 0% | 40% | 100% |
| 0% | 100% | 100% | 0% | 100% |
| 40% | 100% | 60% | 0% | 100% |

[‡], The combination of data sets was bootstrapped 50 times and the training and testing was repeated accordingly.
